# Plasticity manifolds and ion-channel degeneracy govern circadian oscillations of neuronal intrinsic properties in the suprachiasmatic nucleus

**DOI:** 10.1101/2022.07.22.501115

**Authors:** Harshith Nagaraj, Rishikesh Narayanan

## Abstract

**Motivation and methods:** The suprachiasmatic nucleus (SCN) is the master circadian clock of the mammalian brain that sustains a neural code for circadian time through oscillations in the firing rate of constituent neurons. These cell-autonomous oscillations in intrinsic properties are mediated by plasticity in a subset of ion-channels expressed in SCN neurons and are maintained despite widespread neuron-to-neuron variability in ion channel expression profiles. How do SCN neurons undergo stable transitions and maintain precision in intrinsic properties spanning the day-night cycle if several ion channels change concomitantly in a heterogeneous neuronal population? Here, we address this important question using unbiased stochastic searches on the parametric and the plasticity spaces using *populations* of SCN models, each explored for *multiple* valid transitions spanning one complete circadian cycle (day-to-night followed by night-to-day transitions).

**Results:** Our analyses provided three fundamental insights about the impact of heterogeneities on the circadian oscillations of SCN intrinsic properties. First, SCN neurons could achieve signature electrophysiological characteristics (day-like or night-like) despite pronounced heterogeneity in ion channel conductances, with weak pairwise correlations between their conductance values. This *ion-channel degeneracy* precluded the need to maintain precise ionchannel expression profiles for achieving characteristic electrophysiological signatures of SCN neurons, thus allowing for parametric heterogeneities despite functional precision. Second, it was not essential that specific conductances had to change by precise values for obtaining valid day-to-night or night-to-day transitions. This *plasticity degeneracy*, the ability of *disparate combinations of ion-channel plasticity* to yield the same functional transition, confers flexibility on individual neurons to take one of several routes to achieve valid transitions. Finally, we performed nonlinear dimensionality reduction analyses on the valid plasticity spaces and found the manifestation of a low-dimensional *plasticity manifold* in day-to-night transitions, but not in night-to-day transitions. These observations demonstrated that the concomitant changes in multiple ion channels are not arbitrary, but follow a structured plasticity manifold that provides a substrate for stability in achieving stable circadian oscillations.

**Implications:** Our analyses unveil an elegant substrate, involving a synthesis of the *degeneracy* and the *plasticity manifold* frameworks, to effectuate stable circadian oscillations in a heterogeneous population of SCN neurons. Within this framework, the ability of multiple ion channels to change concomitantly provides robustness and flexibility to effectively achieve precise transitions despite widespread heterogeneities in ion-channel expression and plasticity.

## INTRODUCTION

The day-night cycle governs several aspects of the life of organisms on planet Earth. The planetary cycle introduces critical changes to several environmental factors including light levels and temperature, which have differential impacts on different organisms. As these environmental changes set constraints on foraging and other behaviors related to survival, organisms have evolved several adaptation strategies to align their behavior and physiology to the day-night cycle. The suprachiasmatic nucleus (SCN) is the master circadian clock of the mammalian brain that plays a central role in this adaptation, sustaining a neural code for circadian time through the firing rate of constituent neurons. Periodic changes in intrinsic properties are cell-autonomous, maintained by a transcription-translation feedback loop (TTFL). The TTFL is tightly intercoupled with neural activity and several molecular signaling cascades, with the rhythm entrained to the external day-light cycle by retinal projections to the SCN (Reppert and Weaver, 2002; Welsh et al., 2010; Colwell, 2011; Partch et al., 2014; Takahashi, 2017; Hastings et al., 2018; Patton and Hastings, 2018; Harvey et al., 2020).

Cell-autonomous oscillations in neuronal intrinsic properties are mediated by changes in a subset of ion-channels expressed in SCN neurons (Jiang et al., 1997; Itri et al., 2005; Pitts et al., 2006; Belle et al., 2009; Itri et al., 2010; Kudo et al., 2011; Schmutz et al., 2014; Flourakis et al., 2015; Kudo et al., 2015; Paul et al., 2016; Whitt et al., 2018; Harvey et al., 2020; McNally et al., 2020). These circadian oscillations are precisely sustained despite recruiting changes in multiple ion channels and despite pronounced neuron-neuron variability in the intrinsic properties of SCN neurons. The question of how plasticity in these ion channels governs neuronal intrinsic properties through circadian oscillations has remained unexplored, especially considering the high degree of neuron-to-neuron variability observed in SCN neurons. Specifically, how are SCN neurons able to undergo precise circadian transitions in their intrinsic properties despite pronounced variability in the underlying biophysical parameters? Why do so many different ion channels undergo changes during circadian oscillations, when it could have been simpler to design a regulatory mechanism that allows plasticity in a single ion channel subtype? How does the system undergo stable transitions and maintain precision in intrinsic properties spanning the day-night cycle if several ion channels change concomitantly in a heterogeneous neuronal population (Yamada and Forger, 2010; Colwell, 2011; Harvey et al., 2020)?

In this study, we address these important questions by generating a population of day-like SCN model neurons which were biophysically constrained and physiologically validated by several characteristic measurements. We then subjected this day-like model population to an entire cycle of circadian oscillations, not by hand tuning ion-channel plasticity, but by searching for *multiple* valid transitions through an unbiased search of the plasticity space in each model. These unbiased searches were performed independently for the day-to-night and subsequent night-to-day transitions for multiple day-like and night-like models, respectively, together completing one circadian cycle. We constrained the permitted plasticity space for day-to-night and night-to-day transitions by respective biophysical measurements, both in terms of the subset of channels that were allowed to change and the direction in which they could change (Jiang et al., 1997; Itri et al., 2005; Pitts et al., 2006; Belle et al., 2009; Itri et al., 2010; Atkinson et al., 2011; Kudo et al., 2011; Schmutz et al., 2014; Flourakis et al., 2015; Kudo et al., 2015; Paul et al., 2016; Whitt et al., 2018; Harvey et al., 2020; McNally et al., 2020). Together, our approach involved independent and unbiased searches, first involving the parametric space for a valid *population* of day-like models, and subsequently on the plasticity space for *populations* of valid day-to-night and night-to-day transitions.

Our approach involving populations of models and valid transitions allowed for exploring the impact of heterogeneities in both the parametric and plasticity spaces on the circadian oscillations of SCN intrinsic properties. Our analyses provided answers to the questions posed above within the elegant frameworks of degeneracy (Edelman and Gally, 2001; Whitacre and Bender, 2010; Rathour and Narayanan, 2019; Goaillard and Marder, 2021; Kamaleddin, 2022) and plasticity manifolds (Mishra and Narayanan, 2021a). First, we show that SCN neurons could achieve signature electrophysiological characteristics despite pronounced heterogeneity in ion channel conductances and calcium kinetics. It was not essential to maintain biophysical parameters at a precise value for achieving characteristic electrophysiological signatures of SCN neurons under day or night conditions. The parameters showed broad distributions and there were no strong pairwise parametric dependencies, with large pairwise distances between different models in the parametric space. This ion-channel degeneracy was observed in the valid day-like population of SCN models and in the models obtained after valid day-to-night and night-to-day transitions as well.

Second, our approach to look for *multiple* valid transitions from any given SCN model showed that there are several routes to achieve valid day-to-night or night-to-day transitions from any day-like or night-like neuron, respectively. Specifically, it was not essential that specific conductances had to change by precise values for obtaining valid transitions. As disparate combinations of ion-channel plasticity could yield the same functional transition, we refer to this as *plasticity degeneracy* in SCN neurons. Thus, the ability to change multiple ion channels concomitantly allows flexibility by each neuron to take one of several routes to achieve valid transitions, rather than being constrained by changes to one specific ion channel. These analyses also showed that given the inherent heterogeneities in individual neurons, the plasticity routes to achieve valid transitions were dependent on the specific neuron under consideration. Together, the ability of multiple ion channels to change during circadian oscillations provides robustness and flexibility to effectively achieve precise transitions despite widespread heterogeneities in ion-channel expression through plasticity degeneracy.

Finally, we posed the question of how SCN neurons maintained stability through continual transitions if so many ion channels were allowed to change concomitantly. An elegant solution to this problem is to constrain the multi-dimensional plasticity space involving several ion channels to a structured low-dimensional manifold so that individual ion channels do not undergo arbitrary plasticity resulting in instability (Mishra and Narayanan, 2021a). We performed nonlinear dimensionality reduction analyses on the valid day-to-night and night-to-day populations of transitions to explore if valid transitions were constricted within *plasticity manifolds*. Strikingly, we found the manifestation of a low-dimensional manifold in the day-to-night transitions, but not in the night-to-day transitions. These observations demonstrated that the concomitant changes in multiple ion channels are not arbitrary but follow a structure that provides a substrate for stability in achieving valid circadian oscillations. We did not find pairwise correlations between ion-channel plasticity across the 6 channel conductances that underwent plasticity during either transition, thus ruling out the need for a single mechanistic basis for plasticity in all ion-channels.

Together, our analyses involving unbiased stochastic searches on the parametric and the plasticity spaces unveil an elegant substrate, involving a synthesis of the degeneracy and the plasticity manifolds frameworks, to effectuate stable circadian oscillations in a heterogeneous population of SCN neurons. Within this framework, heterogeneities provide a substrate for realizing degeneracy and could emerge from the multiple plasticity routes that can yield valid transitions. Our analyses emphasize the critical need to account for neural-circuit heterogeneities in studying circadian oscillations and provide an overarching framework to assess the emergence and consequences of heterogeneities in the SCN. The manifestation of plasticity manifolds provides a structure that avoids instability by constraining concomitant plasticity to specific combinations of changes that need not necessarily involve correlated changes in all ion channels. These analyses also argue for the SCN as an ideal system to address conceptual and mechanistic questions on the synergy between heterogeneities, long-term plasticity manifolds, and degeneracy in coupled functional emergence across multiple scales.

## METHODS

We employed a single-compartmental cylindrical model of an SCN neuron (Fig. 2*A*; diameter, *d* = 22.3 μm; length, *L* = 22.3 μm). Passive properties were incorporated as an RC circuit, with a specific membrane resistance *R*_m_ and a specific membrane capacitance *C*_m_. *C*_m_ was set at 1 μF/cm^2^ and *R*_m_ was set between 20–40 kΩ.cm^2^ to obtain a membrane time constant (=*R*_m_*C*_m_) of 20-40 ms (Pennartz et al., 1998). The geometry of the neuron was adjusted to obtain a passive input resistance (*R*_m_/*π d L*) in the range of 1.28–2.56 GΩ, matching with electrophysiological measurements from day-like SCN neurons (Hermanstyne et al., 1998; Pennartz et al., 1998). The model comprised 11 active channels: fast sodium (NaF), persistent sodium (NaP), sodium leak (NaLCN), fast delayed rectifier potassium (KFR), slow delayed rectifier potassium (KSR), *A*-type potassium (KA), small-conductance Ca^2+^-activated potassium (SK), big-conductance Ca^2+^-activated potassium (BK), *P*-type calcium (CaP), *L*-type calcium (CaL), and the hyperpolarization-activated cyclic-nucleotide gated nonspecific cationic (HCN) channels.

**Figure 1.**
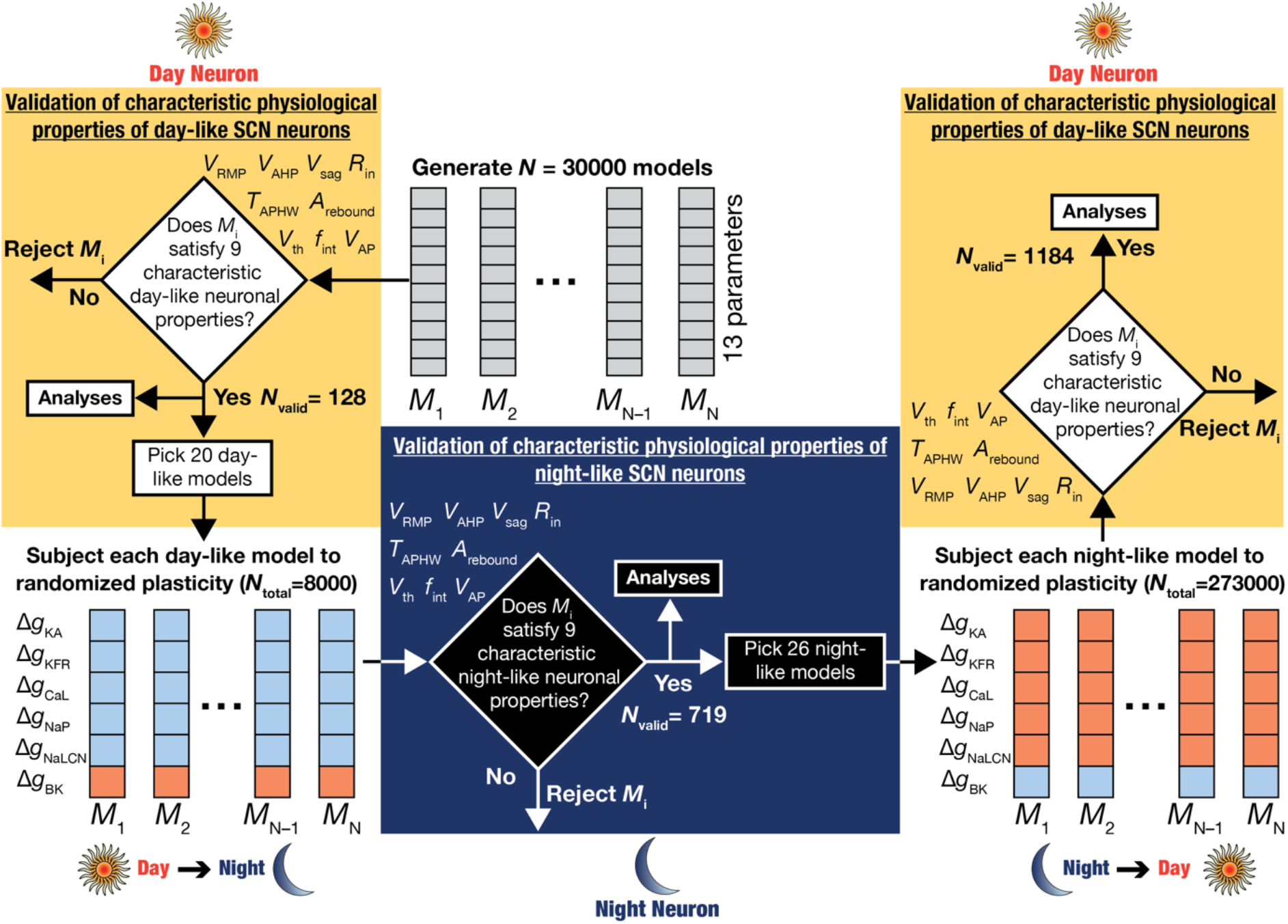
Flowchart demonstrating the overall methodological plan for assessing ion-channel degeneracy and plasticity manifolds in circadian oscillations in SCN neurons. *Left*, The first set of day-like neurons were generated by a *de novo* unbiased search involving 13 different parameters involving 30000 neuron models. Of these, 128 were found to show valid day-like physiological properties. These models were assessed for the expression of ion-channel degeneracy and heterogeneities. As a second step, 20 of these 128 day-like models were picked and subjected to day-to-night transitions that involved plasticity in six different ion channels in electrophysiologically determined directions (red implies increase, blue implies reduction). *Center*, Models subjected to randomized plasticity (*N*_total_=8000 total random transitions) were validated with night-like measurements from SCN neurons, and 719 models derived from the 20 day-like models were found to be valid. These models were assessed for manifestation of ion-channel degeneracy and heterogeneities. The specific combinations of ion-channel plasticity (from respective day-like neurons) that resulted in valid night-like models were subjected to dimensionality reduction analysis to determine the presence of structured plasticity manifolds in the day-to-night transitions. *Right*, As a third step, 26 of these 719 night-like models were picked and subjected to night-to-day transitions that involved plasticity in six different ion channels in electrophysiologically determined directions (red implies increase, blue implies reduction). Models subjected to randomized plasticity (*N*_total_=273000 total random transitions) were validated with day-like measurements from SCN neurons, and 1184 models derived from the 26 night-like models were found to be valid. These models were assessed for manifestation of ion-channel degeneracy and heterogeneities. The specific combinations of ion-channel plasticity (from respective night-like neurons, as well as the original day-like neurons from step 1) that resulted in valid day-like models were subjected to dimensionality reduction analysis to determine the presence of structured plasticity manifolds in these transitions.

**Figure 2.**
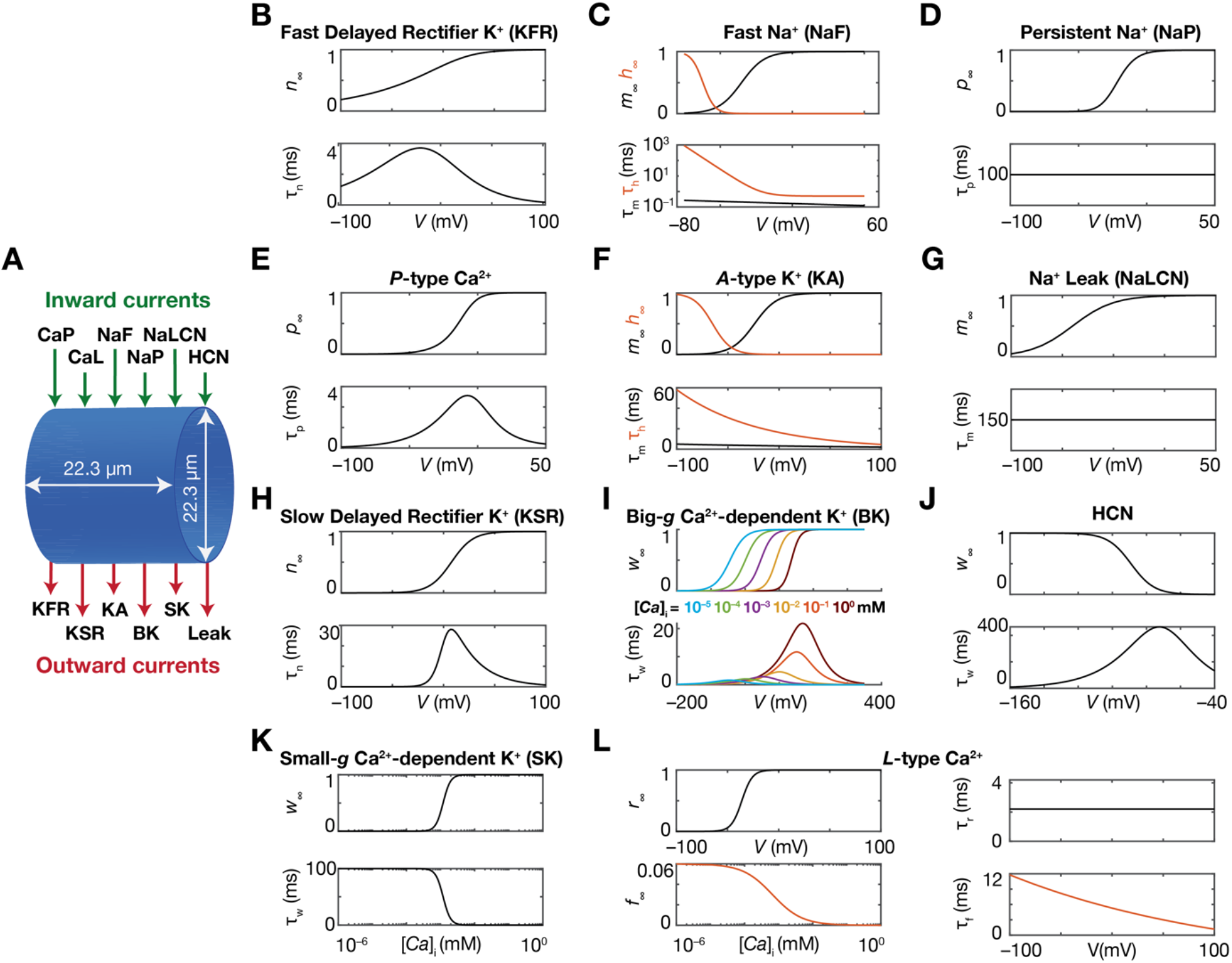
Gating profile and kinetics of the ion channels. (*A*) Representation of the conductance-based model showing the different inward and outward channel currents. (*B–L*) Dependencies of steady-state values and the kinetics of the gating variables associated with the different ion channel conductances in the model. In panels C, F, and L, black and orange lines represent activation and inactivation gating variables, respectively.

### Ion-channel gating properties and kinetics

The channel kinetics and voltage-dependencies for KFR and KA were adopted from (Bouskila and Dudek, 1995); for SK and CaP from (Huang et al., 2012); for NaF from (Sim and Forger, 2007); for NaP from (Paul et al., 2016); for NaLCN from (Chua et al., 2020); for BK from (Cui et al., 1997); for HCN from (de Jeu and Pennartz, 1997); for CaL from (Diekman et al., 2013) and KSR from (Itri et al., 2005).

All channel models were based on the Hodgkin-Huxley formulation (Hodgkin and Huxley, 1952). The sodium, potassium and HCN channels followed the Ohmic formulation, while the calcium channels followed the Goldman-Hodgkin-Katz (GHK) formulation (Goldman, 1943; Hodgkin and Katz, 1949). The reversal potentials for Na^+^, K^+^, and HCN were set at +45, −97 and −30 mV respectively (de Jeu and Pennartz, 1997; Sim and Forger, 2007). The evolution of cytosolic calcium concentration [*Ca*]_*i*_ was dependent on the current through the voltage-gated calcium channels and involved a first-order decay with a decay time constant in the range of 1750-2240 ms (Diekman et al., 2013):

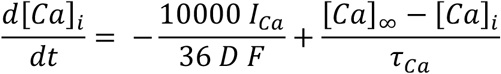

where *F* represents Faraday’s constant, *τ_Ca_* represents the decay time constant of calcium in SCN neurons, *D* was the depth of the shell into which calcium influx occurs (*D* = 0.1 μm), and [*Ca*]_∞_ was the steady state value of [*Ca*]_*i*_ set at 50 nM.

Channel models were directly adopted from previous studies when available. In case channel models were not available, they were defined to match respective electrophysiological measurements. The channels were described by either one or two gating particles, with each particle following the first-order kinetics:

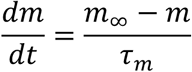

where *m*_∞_ and *τ*_m_ defined the steady-state value and the time constant of the state variable governing the gating particle, respectively. Channel gating and kinetics were adjusted to account for temperature dependence as well. The details of the gating properties as well as kinetics for each channel are as follows (Fig. 2).

#### Fast Delayed Rectifier Potassium Channel (KFR)

The model for KFR channel was obtained by fitting the corresponding electrophysiological data from (Bouskila and Dudek, 1995), and the current through this channel was as follows (Fig. 2*B*):

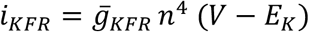

The activation gating particle was governed by:

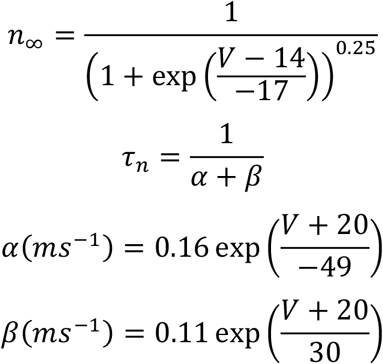

#### Fast Sodium Channel (NaF)

The model for NaF was adopted from (Sim and Forger, 2007), and the current through this channel was (Fig. 2*C*):

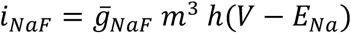

The activation gating variable was defined by:

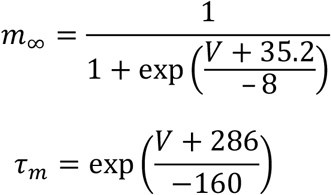

The inactivation gating particle was defined by:

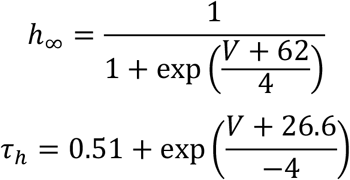

#### Persistent Sodium Channel (NaP)

The model for NaP was adopted from (Paul et al., 2016). The current through the channel was given by (Fig. 2*D*):

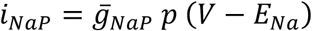

The activation gating variable was governed by:

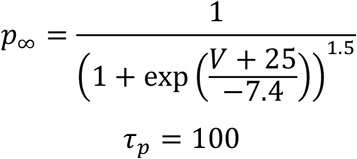

#### P-type Calcium Channel (CaP)

The model for CaP (Fig. 2*E*) was adapted from (Huang et al., 2012). The current through this channel followed GHK conventions. The default extracellular and cytosolic calcium concentrations were set at 2 mM and 100 nM respectively.

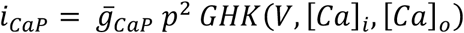

The activation gating variable was governed by (Fig. 2*E*):

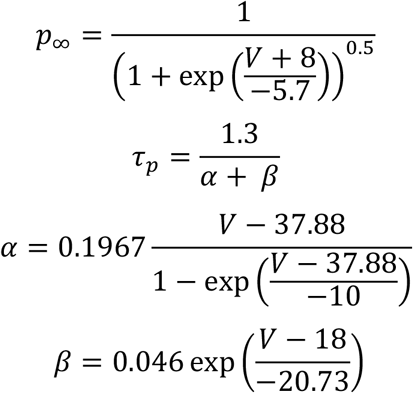

#### A-type Potassium Channel (KA)

The model for KA was obtained by fitting the corresponding electrophysiological data from (Bouskila and Dudek, 1995). The current through the channel was defined as follows (Fig. 2*F*):

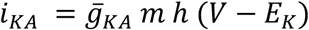

The activation gating variable was described by:

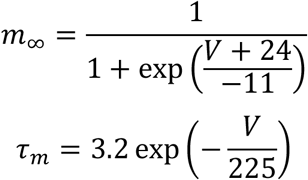

The inactivation gating variable was described by:

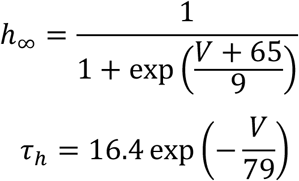

#### Sodium Leak Channel (NaLCN)

The model for NaLCN was obtained by fitting the corresponding electrophysiological data from (Chua et al., 2020). The current through the channel was given by (Fig. 2*G*):

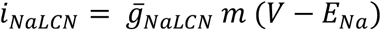

The activation gating variable was defined by:

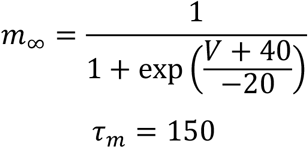

#### Slow Delayed Rectifier Potassium Channel (KSR)

The model for KSR was obtained by fitting the corresponding electrophysiological data from (Itri et al., 2005). The current through this channel was as follows (Fig. 2*H*):

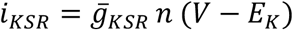

The activation gating particle was governed by:

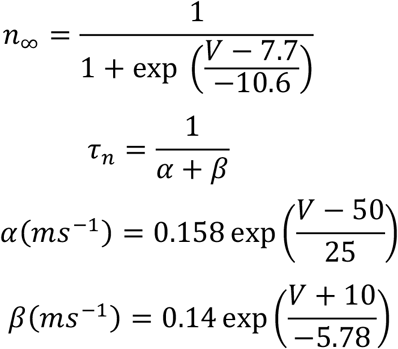

#### Big-Conductance Ca^2+^-Activated Potassium Channel (BK)

The model for BK was obtained by fitting the corresponding electrophysiological data from (Cui et al., 1997). The current through the channel described by:

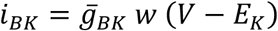

The activation gating variable was governed by (Fig. 2*I*):

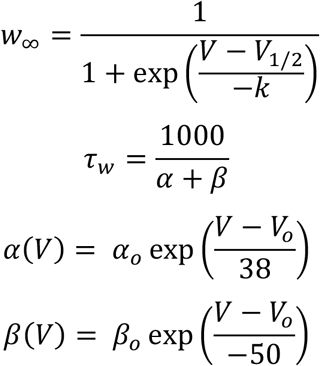

The [*Ca*]_*i*_ dependencies of gating kinetics were accounted for:

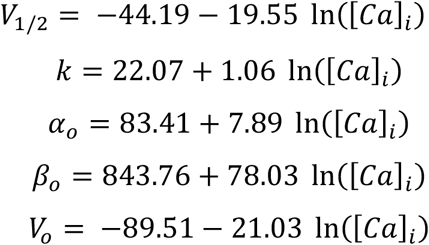

#### Hyperpolarization-activated cyclic-nucleotide gated channel (HCN)

The model for the HCN channel was obtained by fitting the corresponding electrophysiological data from (de Jeu and Pennartz, 1997). The current through the channel was defined as follows (Fig. 2*J*):

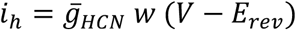

The activation gating variable of the channel was described by:

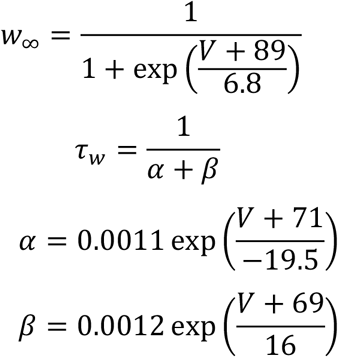

#### Small-Conductance Ca^2+^-Activated Potassium Channel (SK)

The model for SK channel was adapted from (Huang et al., 2012), with the current through the channel given by:

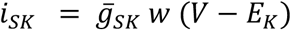

The activation gating variable was described by (Fig. 2*K*):

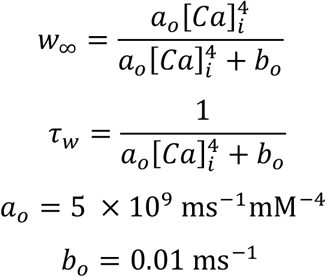

where [*Ca*]_*i*_ was specified in mM.

#### L-type Calcium Channel (CaL)

The model for CaL was adapted from (Diekman et al., 2013). The current through this channel followed GHK conventions. The default extracellular and cytosolic calcium concentrations were set at 2 mM and 100 nM respectively.

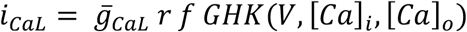

The activation gating variable was governed by (Fig. 2*L*):

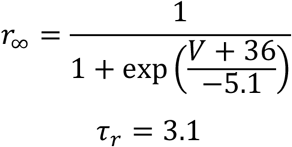

The inactivation gating variable, dependent on calcium and voltage, was defined as:

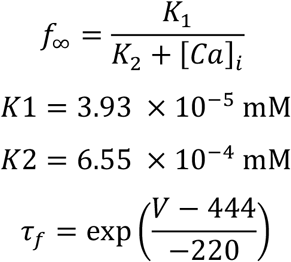

### Measurements

#### Resting Membrane Potential and Intrinsic Firing Frequency

The resting membrane potential (*V*_R*MP*_), in the absence of any active conductance was set at – 65 mV. The active currents were allowed to interact for 2 s to allow the membrane potential to settle down into a steady state, which may be spontaneously firing or silent. All measurements were obtained after this initial period of 2 s when the neuron’s RMP reached a steady-state value. Since most day-like neurons manifest spontaneous action potential firing, *V*_R*MP*_ was measured by median filtering the voltage traces over a period of 5 s and computing the mean of the filtered trace (Fig. 3*A*). The intrinsic frequency (*f*_int_) was computed as the total number of spikes over a period of 5 s.

**Figure 3.**
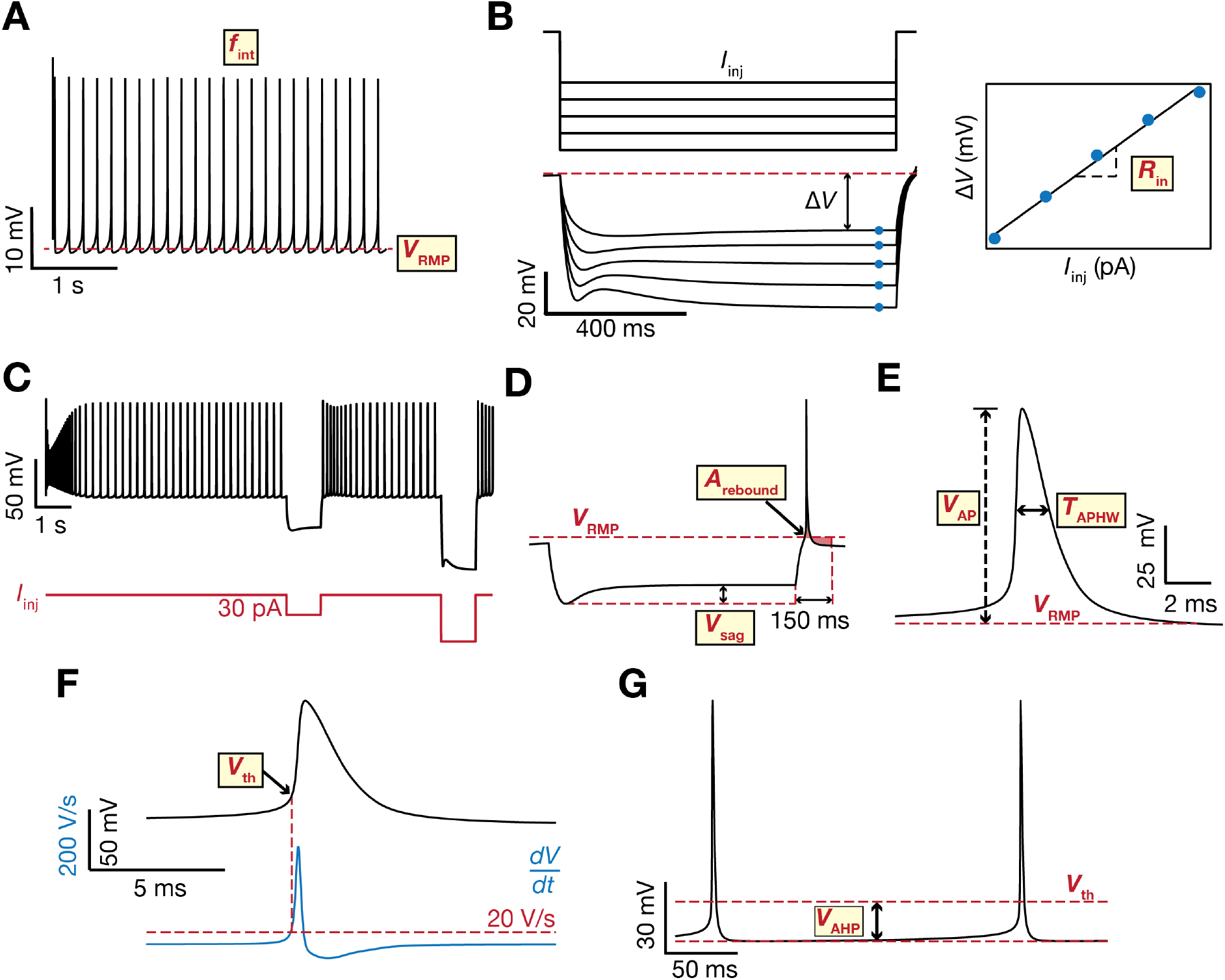
Electrophysiological measurements used to characterize and validate the models. (*A*) The resting membrane potential (*V*_RMP_, red line) was measured as the mean of the median-filtered voltage trace computed over a duration of 5 s. (*B*) The input resistance (*R*_in_) was measured as the slope of the linear fit to the steady state voltage deflection obtained by injecting 5 hyperpolarizing current pulses (–30 pA to –70 pA, in steps of 10 pA) for a period of 1 s. (*C*) Voltage response of an example neuron (black) to current injections (red). The neuron is spontaneously active, with a characteristic sag and rebound depolarization seen due to the injection of hyperpolarizing current pulses. (*D*) The sag (*V*_sag_) and rebound depolarization (*V*_rebound_) were measured after injecting a hyperpolarizing current pulse of –30 pA for a duration of 1 s. The sag was the difference between the steady state voltage and the minimum voltage obtained after injecting the current pulse. Rebound depolarization was the area under the voltage-time curve (shaded in red) over a period of 150 ms after the current injection. (*E*) The spike amplitude (*V*_AP_) was quantified as the difference between the maximum voltage attained by the spike with reference to *V*_RMP_. Spike width (*T*_APHW_) was quantified as the difference between the two timepoints when the voltage attained was half the spike amplitude with reference to *V*_RMP_. (*F*) The spike threshold (*V*_th_) was measured as the voltage attained at the point when the rate of change of voltage with time (*dV*/*dt*, blue) exceeded 20 V/s. (*G*) The spike after-hyperpolarisation (*V*_AHP_) was quantified as the difference between the minimum voltage attained within 40 ms after the spike attained its maximum value (lower red line) and *V*_th_.

#### Input Resistance

The input resistance (*R_in_*) was computed from the steady state voltage response to 5 hyperpolarizing current pulses between −30 pA to −70 pA, in steps of 10 pA, injected for a duration of 1 s. The steady-state voltages were plotted against the corresponding amplitude of injected current, and the slope of the linear fit to this model was taken as *R*_in_ (Fig. 3*B*).

#### Sag and Rebound Depolarization

A hyperpolarizing pulse of −30 pA was injected for a duration of 1 s (Fig. 3C–*D*) to measure the sag (*V_sag_*) and area under rebound depolarization (*A*_rebound_). *V_sag_* was measured as the difference between the peak deflection (*V_peak_*) attained following injection of the pulse and the steady state value of deflection (*V_ss_*). *A*_rebound_ was measured as the area under the voltage-curve, with reference to *V_RMP_*, for a period of 150 ms following termination of hyperpolarization.

#### Spike Properties

Spike properties were measured from the first action potential elicited after an initial period of 2 s. The spike amplitude (*V_AP_*) was computed as the difference between the peak voltage attained by the spike and *V*_R*MP*_ (Fig. 3*E*). The spike width (*T*_APHW_) was measured as the difference between the two time points at which the difference between the voltage and resting membrane potential was half of the spike amplitude (Fig. 3*E*). The spike threshold (*V*_th_) was measured as the voltage at the point at which the rate of change of voltage exceeded 20 V/s (Fig. 3*F*). The amplitude of the spike after-hyperpolarization (*V*_AHP_) was computed as the difference between the minimum voltage attained after the spike and *V*_th_ (Fig. 3*G*).

### Multi-Parametric Multi-Objective Stochastic Search

To arrive at a heterogeneous population of day-like SCN neurons and to probe the manifestation of ion-channel degeneracy in the neuronal phenotypes, a multiparametric, multiobjective stochastic search (MPMOSS) algorithm was employed (Foster et al., 1993; Prinz et al., 2003; Rathour and Narayanan, 2012, 2014; Basak and Narayanan, 2018a; Mittal and Narayanan, 2018; Mishra and Narayanan, 2019; Jain and Narayanan, 2020). Here, a stochastic search was performed over 13 parameters (Table 1) and the models obtained were validated using the 9 supra- and subthreshold measurements (Table 2) based on the *in vitro* electrophysiological bounds (Hermanstyne et al., 1998; Pennartz et al., 1998; Atkinson et al., 2011). As part of the MPMOSS algorithm, each parameter was randomly picked from a uniform distribution (Table 1). From this model, the 9 intrinsic measurements were computed and validated against the respective day-like electrophysiological bounds (Table 2). A model that satisfied all the 9 criteria for validation was used for further analysis. This process was repeated for 30,000 such unique randomized picks spanning the 13 parameters (Fig. 1). Parameters were allowed to assume arbitrary values within their respective bounds, thereby avoiding artificial discretization of the parametric space.

**Table 1.** Range of Parameters used in generating the day-like model population.

|  | Parameter, symbol (Unit) | Lower bound | Upper bound |
| --- | --- | --- | --- |
| 1 | Maximal conductance of KFR channels, $\bar{g}_{\text{KFR}}$ (mS/cm<sup>2</sup>) | 0.1 | 1 |
| 2 | Maximal conductance of KSR channels, $\bar{g}_{\text{KSR}}$ (mS/cm<sup>2</sup>) | 0.1 | 1 |
| 3 | Maximal conductance of KA channels, $\bar{g}_{\text{KA}}$ (mS/cm<sup>2</sup>) | 0.01 | 0.1 |
| 4 | Maximal conductance of NaF channels, $\bar{g}_{\text{NaF}}$ (S/cm<sup>2</sup>) | 0.05 | 0.5 |
| 5 | Maximal conductance of NaP channels, $\bar{g}_{\text{NaP}}$ (mS/cm<sup>2</sup>) | 0.01 | 0.1 |
| 6 | Maximal conductance of CaL channels, $\bar{g}_{\text{CaL}}$ (mS/cm<sup>2</sup>) | 0.1 | 1 |
| 7 | Maximal conductance of CaP channels, $\bar{g}_{\text{CaP}}$ (mS/cm<sup>2</sup>) | 0.1 | 1 |
| 8 | Maximal conductance of SK channels, $\bar{g}_{\text{SK}}$ (μS/cm<sup>2</sup>) | 1 | 10 |
| 9 | Maximal conductance of BK channels, $\bar{g}_{\text{BK}}$ (S/cm<sup>2</sup>) | 0.01 | 0.1 |
| 10 | Maximal conductance of HCN channels, $\bar{g}_{\text{HCN}}$ (μS/cm<sup>2</sup>) | 5 | 50 |
| 11 | Maximal conductance of NaLCN channels, $\bar{g}_{\text{NaLCN}}$ (μS/cm<sup>2</sup>) | 5 | 50 |
| 12 | Specific membrane resistance, $R_m$ (kΩ.cm<sup>2</sup>) | 20 | 40 |
| 13 | Calcium decay time constant, $\tau_{\text{Ca}}$ (s) | 1.75 | 2.24 |

**Table 2.** Range of measurement bounds used for validating the day-like and night-like neurons.

|  | Measurement (Unit) | Day neurons |  | Night neurons |  |
| --- | --- | --- | --- | --- | --- |
|  |  | Lower bound | Upper bound | Lower bound | Upper bound |
| 1 | $V_{\text{RMP}}$ (mV) | -60 | -52 | -75 | -65 |
| 2 | $R_{\text{in}}$ (GΩ) | 0.768 | 1.812 | 0.5 | 1.5 |
| 3 | $V_{\text{AP}}$ (mV) | 70 | - | 70 | - |
| 4 | $V_{\text{th}}$ (mV) | -44.7 | -36.5 | - | - |
| 5 | $T_{\text{APHW}}$ (ms) | 1 | 2 | 1 | 2 |
| 6 | $V_{\text{AHP}}$ (mV) | -25.8 | -16.8 | -31 | -17 |
| 7 | $A_{\text{rebound}}$ (mV.ms) | -510 | 966 | -510 | 966 |
| 8 | $f_{\text{int}}$ (Hz) | 3 | 7 | 0 | 2 |
| 9 | $V_{\text{sag}}$ (mV) | 2 | 10 | 2 | 16 |

### Day to Night Transitions

To examine plasticity of individual ion channels during day to night transitions, night models were generated by picking day-like neurons from the valid model population (Fig. 1). Consistent with electrophysiological observations on which channels change in what direction during the circadian oscillations (Itri et al., 2005; Pitts et al., 2006; Itri et al., 2010; Atkinson et al., 2011; Flourakis et al., 2015; Paul et al., 2016; Harvey et al., 2020; McNally et al., 2020), day-to-night transitions in these models was implemented by changing 6 parameters: the conductances of the KFR, KA, NaP, CaL, BK and NaLCN channels. While a reduction in the night conductance compared to the day conductance has been observed experimentally in case of the KFR, KA, NaP, CaL and NaLCN channels, an increase in night conductance relative to the day conductance has been observed in case of the BK channel. The signs of these experimentally observed day-to-night transition-induced changes were enforced in our model (Fig. 1).

The search for transitions that would produce night-like SCN neurons from their day-like counterparts was implemented using an MPMOSS algorithm. The parametric space for the stochastic search algorithm contained sign-enforced changes in the six conductance values (Table 3) representing the transition. While the range of the distribution for ion channels that showed a reduction was restricted to (−1,0) (values higher than 1 would lead negative conductance values, and 0 represents no change), it was (0,10) in case of ion channels that would show an increase, to span a broad parametric space (Table 3). For a given model, in each iteration, a randomized percentage change was picked for each of the six conductance values within their respective bounds (Table 3). These changes were then introduced into the specific day-like model neuron, and the 9 characteristic electrophysiological properties were measured. If the neuron matches all 9 night-like properties (Table 2), the randomized transition was declared a valid transition and the model that the transition yielded was called as a valid night-like model (Fig. 1). This process was repeated for each of the different day-like neurons for several iterations to generate several valid night-like neurons from each day-like neuron. The parameters of valid night-like neurons were used for assessing ion-channel degeneracy, and the valid day-to-night transitions were used to investigate the manifestation of plasticity manifolds using different nonlinear dimensionality reduction techniques.

**Table 3.** Range of plasticity parameters used in the day-to-night transitions.

| | Parameter (Unit) | Range of $\Delta g$ | Relation between $g_{day}$ and $g_{night}$ |
| --- | --- | --- | --- |
| 1 | $\bar{g}_{KFR}$ (mS/cm<sup>2</sup>) | (0, 1) | $g_{Night}^{KFR} = g_{Day}^{KFR} (1 - \Delta g)$ |
| 2 | $\bar{g}_{KA}$ (mS/cm<sup>2</sup>) | (0, 1) | $g_{Night}^{KA} = g_{Day}^{KA} (1 - \Delta g)$ |
| 3 | $\bar{g}_{NaP}$ (mS/cm<sup>2</sup>) | (0, 1) | $g_{Night}^{NaP} = g_{Day}^{NaP} (1 - \Delta g)$ |
| 4 | $\bar{g}_{CaL}$ (mS/cm<sup>2</sup>) | (0, 1) | $g_{Night}^{CaL} = g_{Day}^{CaL} (1 - \Delta g)$ |
| 5 | $\bar{g}_{BK}$ (S/cm<sup>2</sup>) | (0, 10) | $g_{Night}^{BK} = g_{Day}^{BK} (1 + \Delta g)$ |
| 6 | $\bar{g}_{NaLCN}$ (μS/cm<sup>2</sup>) | (0, 1) | $g_{Night}^{NaLCN} = g_{Day}^{NaLCN} (1 - \Delta g)$ |

### Night to Day Transitions

An independent MPMOSS algorithm was used to arrive at valid night-to-day transitions from night-like neurons and resultant valid day-like neurons (Fig. 1). The procedure and the ion-channel conductances were identical to day-to-night transitions, with the sign of transitions in individual transitions reversed for night-to-day transitions (Table 4) compared to their day-to-night counterparts. As before, the stochastic search spanned the plasticity space involving these 6 conductance values on different night-like neurons. The validation of these randomly generated models (derived from randomized transitions on different night-like neurons) was against day-like physiological properties (Table 2). Neurons that satisfied all 9 day-like properties were declared valid and the transitions that resulted in these valid day-like models were declared valid night-to-day transitions. The parametric space of valid models and associated transitions were then used to explore the manifestation of ion-channel degeneracy and plasticity manifolds.

**Table 4.** Range of plasticity parameters used in the night-to-day transitions.

| | Parameter (Unit) | Range of $\Delta g$ | Relation between $g_{day}$ and $g_{night}$ |
| --- | --- | --- | --- |
| 1 | $\bar{g}_{KFR}$ (mS/cm<sup>2</sup>) | (0, 10) | $g_{Day}^{KFR} = g_{Night}^{KFR} (1 + \Delta g)$ |
| 2 | $\bar{g}_{KA}$ (mS/cm<sup>2</sup>) | (0, 10) | $g_{Day}^{KA} = g_{Night}^{KA} (1 + \Delta g)$ |
| 3 | $\bar{g}_{NaP}$ (mS/cm<sup>2</sup>) | (0, 10) | $g_{Day}^{NaP} = g_{Night}^{NaP} (1 + \Delta g)$ |
| 4 | $\bar{g}_{CaL}$ (mS/cm<sup>2</sup>) | (0, 10) | $g_{Day}^{CaL} = g_{Night}^{CaL} (1 + \Delta g)$ |
| 5 | $\bar{g}_{BK}$ (S/cm<sup>2</sup>) | (0, 1) | $g_{Day}^{BK} = g_{Night}^{BK} (1 - \Delta g)$ |
| 6 | $\bar{g}_{NaLCN}$ (μS/cm<sup>2</sup>) | (0, 10) | $g_{Day}^{NaLCN} = g_{Night}^{NaLCN} (1 + \Delta g)$ |

### Computational Details

All simulations were performed using NEURON programming environment (Carnevale and Hines, 2006) with an integration step size of 25 μs. All data analyses and plotting were done using custom-written scripts within IGOR Pro (Wavemetrics) and MATLAB (Mathworks) environments.

## RESULTS

The goal of this study was to assess constraints on ion channels and intrinsic properties in the emergence of SCN neurons endowed with characteristic physiological properties and signature circadian oscillations in their intrinsic physiology (Fig. 1). As the use of single hand-tuned models does not capture the heterogeneity that is characteristic of all biological systems, we generated a population of neurons that were endowed with characteristic biophysical properties (Fig. 2) and satisfied signature electrophysiological properties of SCN neurons (Fig. 3). We used this population of neurons to assess heterogeneity of measurements, degeneracy in parametric space, and the evolution of the parametric and measurement space as the population was subjected to day-to-night followed by night-to-day transitions (Fig. 1). The overall goal was to explore heterogeneities in and constraints on ion channels and their plasticity in the emergence of SCN neurons and in accomplishing the signature circadian transitions over a full cycle (day-to-night and night-to-day; Fig. 1).

### Ion-channel degeneracy in SCN neurons manifesting day-like characteristics

We built a single-compartmental, conductance-based model of an SCN neuron, incorporating 12 active and passive conductances with characteristics derived from biophysical measurements from SCN neurons (Fig. 2). An unbiased stochastic search algorithm was employed spanning 13 parametric values, which accounted for these channel conductances as well as calcium decay (Table 1). A randomized population of 30,000 unique model neurons was generated by sampling this 13-dimensional parametric space and a set of 9 electrophysiological measurements were recorded from each model (Fig. 3). These measurements were validated by comparing model measurements against established experimental bounds characteristic of day-like SCN neurons (Table 2). A small subset of 128 models (128/30000=0.4%) satisfied all 9 day-like measurement constraints (Table 2) and were declared as valid day-like SCN neurons.

These valid models manifested heterogeneous physiological properties within the valid electrophysiological bounds (Fig. 4*A*), thus providing us with a heterogeneous population of valid day-like SCN models reflective of heterogeneities observed in electrophysiological recordings. We asked if the different measurements associated with the 128 valid models manifested pairwise correlation. We found that most of these pairwise correlations were weak (Fig. 4*B*), implying that these 9 measurements were characterizing distinct aspects of SCN physiology. The few strong pairwise correlations observed were expected because of their cross-dependencies arising either from the inherent nature of measurement or because of the common set of ion channels that are expected to govern these properties in individual neurons. For instance, action potential threshold, resting membrane potential, AHP potential, and action potential amplitude showed strong correlations because of how these measurements are performed in a spontaneously firing neuron (Fig. 3). The firing rate *f*_int_ was expectedly higher in neurons with depolarized RMP, manifesting as a strong positive correlation (Fig. 4*B*).

**Figure 4.**
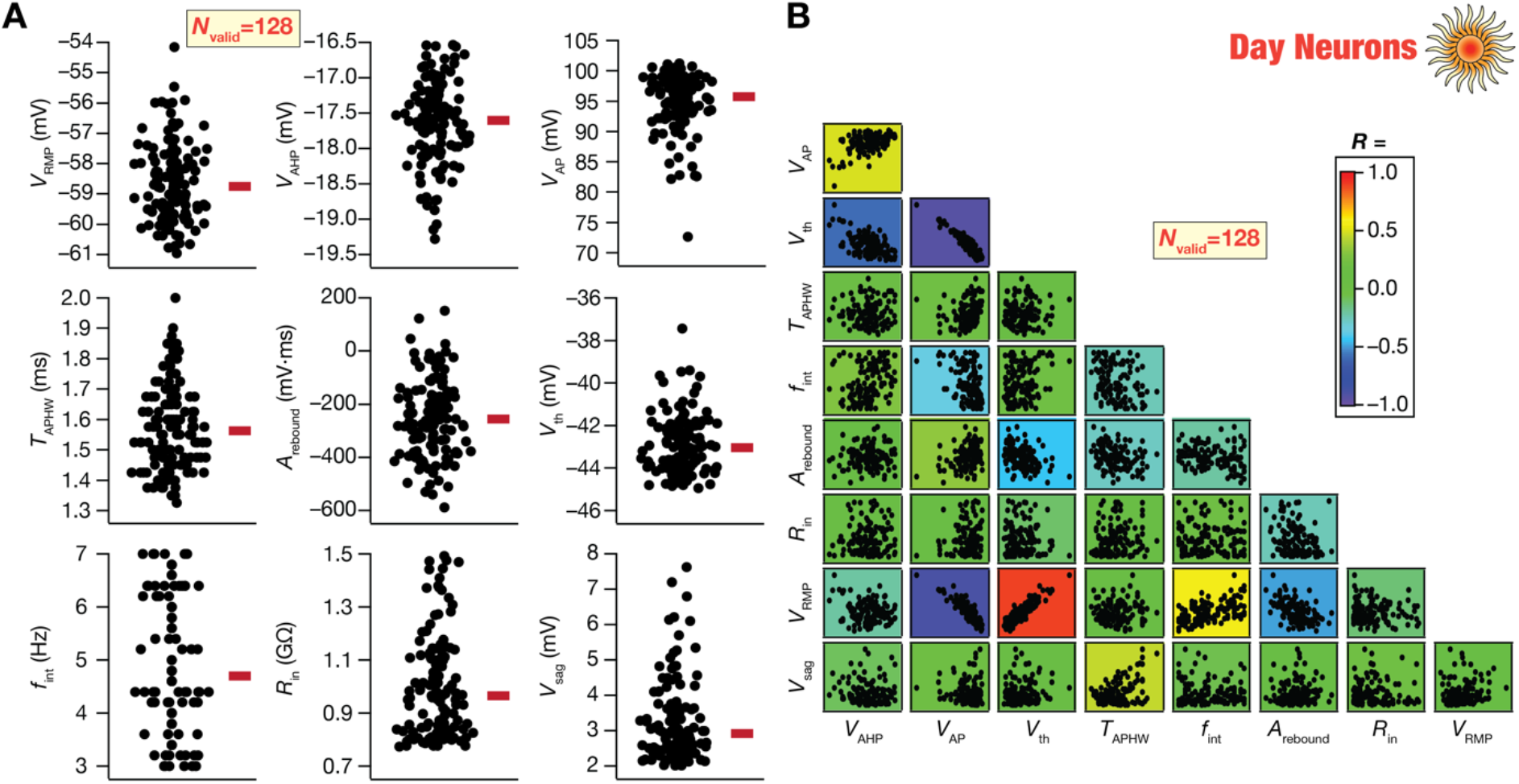
Heterogenous distribution of measurements from day-like SCN neurons. (*A*) Bee-swarm plots of measurements from 128 different SCN neurons exhibiting day-like phenotype. Red bars represent the median values. The neurons were obtained using the MPMOSS algorithm and were validated using the respective electrophysiological bounds (Table 1). The 128 models shown here were obtained from 30,000 runs of the MPMOSS algorithm and exhibited pronounced heterogeneities within their respective bounds (*Table 1*). (*B*) Pairwise correlations between the different measurements used to characterize and validate day-like neuronal models. The scatter plots are overlaid on top of the color-coded scatter plot matrix showing the respective Pearson’s correlation coefficient values.

How constrained were the parametric distributions in these models that exhibited characteristic signatures of day-like SCN neurons? Were they clustered around a specific location or were they distributed across a large swath of the allowed range (Table 1)? To address these, we first picked 7 models that showed similar physiological measurements (Fig. 5*A–F*). We found that the parametric distributions of these models showing similar functions spanned their entire ranges (Fig. 5*G*), thus providing a line of evidence on the manifestation of ion-channel degeneracy in these models. Specifically, these illustrative examples provide evidence that it is not essential to maintain the conductances of different channels at specific levels for achieving the characteristic physiological properties of day-like SCN neurons. We assessed the distributions of parameters across all 128 valid models and found them to show widespread distributions spanning the entire stretch of their respective parametric ranges (Fig. 6*A*, histograms; Table 1).

**Figure 5.**
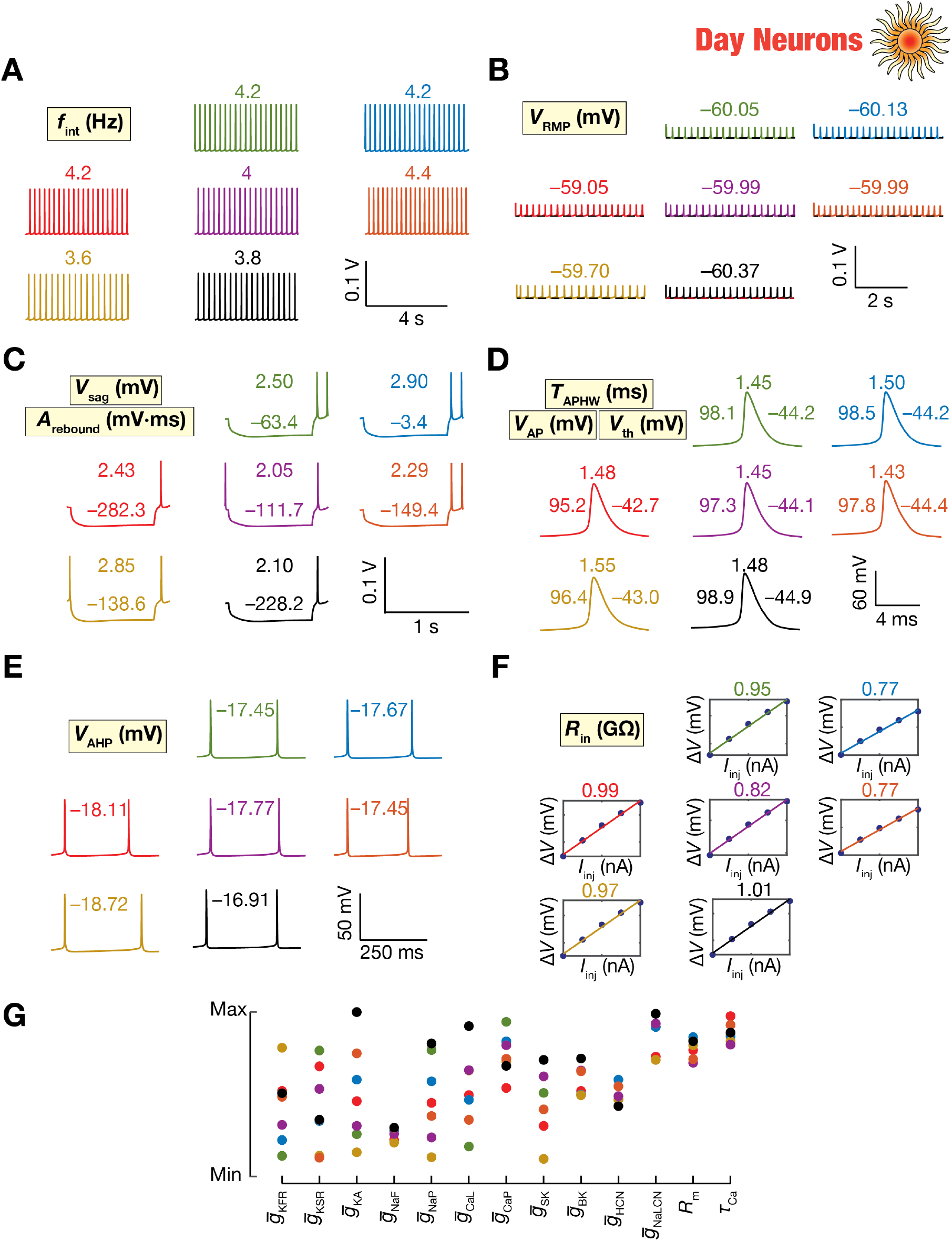
Parametric degeneracy among day-like SCN neurons showing similar characteristic physiological properties. (*A–F*) Measurements of 7 different day-like SCN neurons (picked from the 128 shown in Fig. 4) which exhibit similar characteristic physiology. Different colors represent different neurons, and the values of the corresponding measurements are shown above the traces and were similar across the 7 neurons. (*G*) The 7 models showed pronounced heterogeneity in their parametric values, with no clustering in parameters, providing an illustration of the expression of degeneracy.

**Figure 6.**
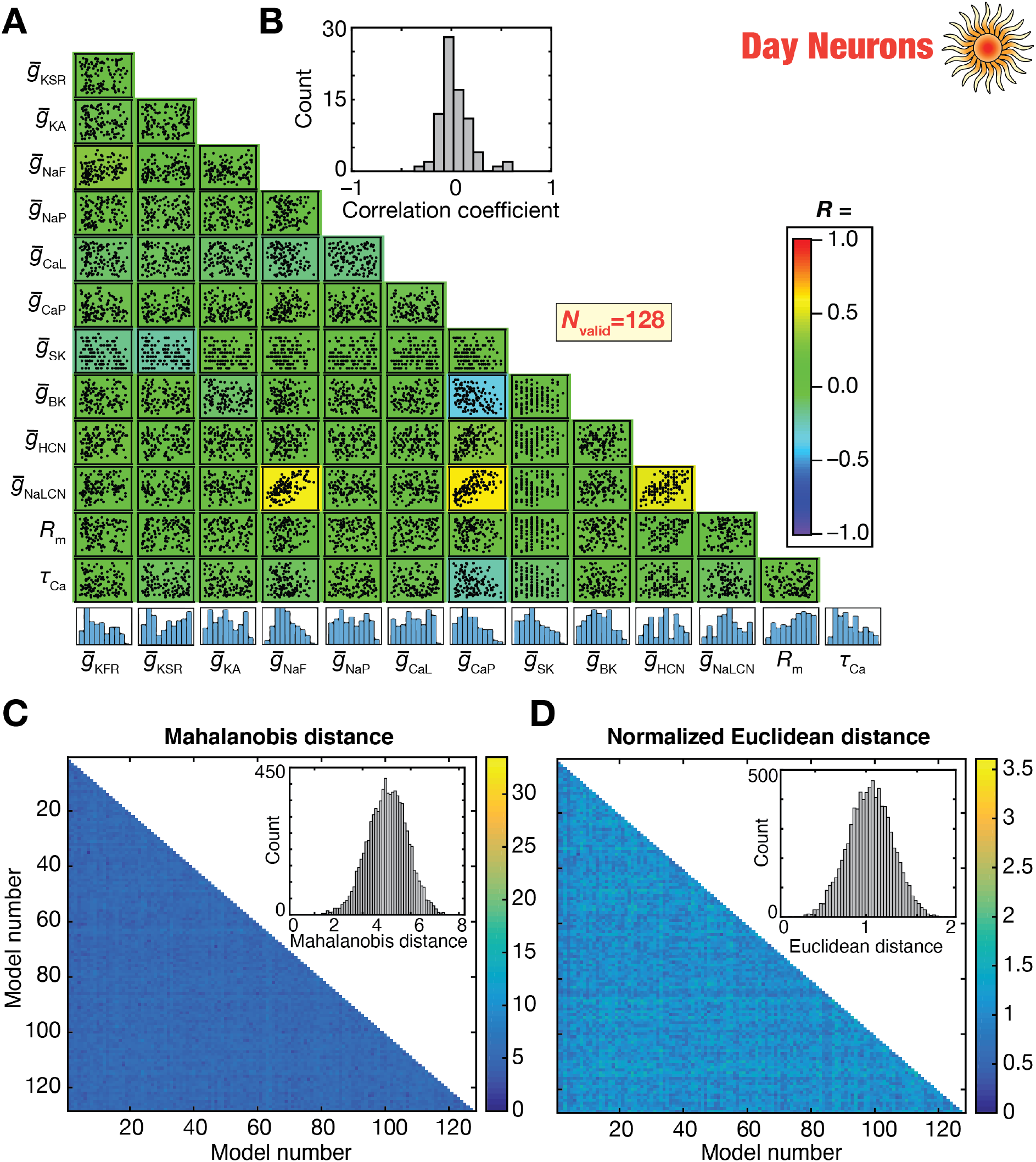
The day-like neuronal model population manifested parametric heterogeneity and weak pairwise correlations in the parametric space. (*A*) Scatter-plot matrix between the different parametric values of the 128 day-like SCN models that are shown in Fig. 4. The background color represents the value of the Pearson correlation coefficient between the different parameters. Histograms representing the distributions of individual parameters is shown on the bottom-most row. (*B*) Histogram representing the distribution of the correlation values shown in panel A. (*C–D*) Heterogeneities in model parameters quantified with Mahalanobis (*C*) or normalized Euclidean (*D*) distances. The matrices represent the pairwise distance between the parametric vectors defining the 128 models. Insets show the histogram of the values in the distance matrix.

How did models achieve characteristic physiology despite such widespread distribution of underlying parameters? Was variability in the expression of one ion channel subtype compensated by specific differences in another conductance value? To address this, we computed pairwise correlations between the different parameters across the 128 valid models (Fig. 6*A–B*). We found most pairwise correlations to be weak (Fig. 6*A–B*), suggesting that variability in one channel parameter was not compensated by variability in another, but because of synergistic interactions between several ion channels defining these model populations. Importantly, we also found that pairwise distances between these models in the 13-dimensional space, computed either through Mahalanobis distance (Fig. 6*C*) or normalized Euclidean (Fig. 6*D*) distance, showed large distances between these models, thus ruling out clustering of these models in the parametric space. The results of linear dimensionality reduction analysis on the measurement and parametric spaces of these 128 neurons, using principal component analysis (PCA), did not show any clustering (Fig. S1). The widespread distribution of model parameters or weak pairwise correlations or large pairwise distances does not imply that any arbitrary combination of these 13 parameters can yield valid day-like SCN model neurons. A vast majority (29872/30000=99.57%) of the parametric combinations spanning this exact same range were in fact rejected during the validation process yielding only 128 valid models out of the 30000 random samples tested. Thus, these observations imply that there are disparate, yet *specific combinations* of parameters, which do not show discernable structure in pair-wise relationships or in individual distributions, that can yield similar characteristic physiological properties observed in SCN neurons. In other words, *specific combinations* of disparate structural components (ion channel subtypes) governed by a global structure in the parametric space yielded similar functions, a phenomenon that has traditionally been defined as degeneracy in biological systems (Edelman and Gally, 2001; Rathour and Narayanan, 2019; Goaillard and Marder, 2021). Together, our analyses involving a heterogeneous population of day-like SCN neurons manifesting widespread parametric variability provide evidence for the manifestation of ion-channel degeneracy in the concomitant emergence of several characteristic properties of SCN neurons.

### Plasticity degeneracy in achieving valid day-to-night transitions unveiled ion-channel degeneracy in night-like neurons

A well-studied signature characteristic of SCN neurons is their ability to undergo circadian modulation in their intrinsic properties, manifesting different electrophysiological characteristics during day *vs*. night periods. These changes in electrophysiological properties are mediated by changes in a specific subset of ion channels, with channel-specific directions in such changes observed during night-to-day and day-to-night transitions. Specifically, there are lines of evidence for reductions in KA, KFR, CaL, NaP, and NaLCN channel conductances and a concomitant increase in BK conductance during day-to-night transitions (Itri et al., 2005; Pitts et al., 2006; Itri et al., 2010; Flourakis et al., 2015; Paul et al., 2016; Whitt et al., 2018; McNally et al., 2020). The direction of changes of this subset of channels is opposite for the night-to-day transitions, with the cycle repeated over the circadian period (Fig. 7*A*).

**Figure 7.**
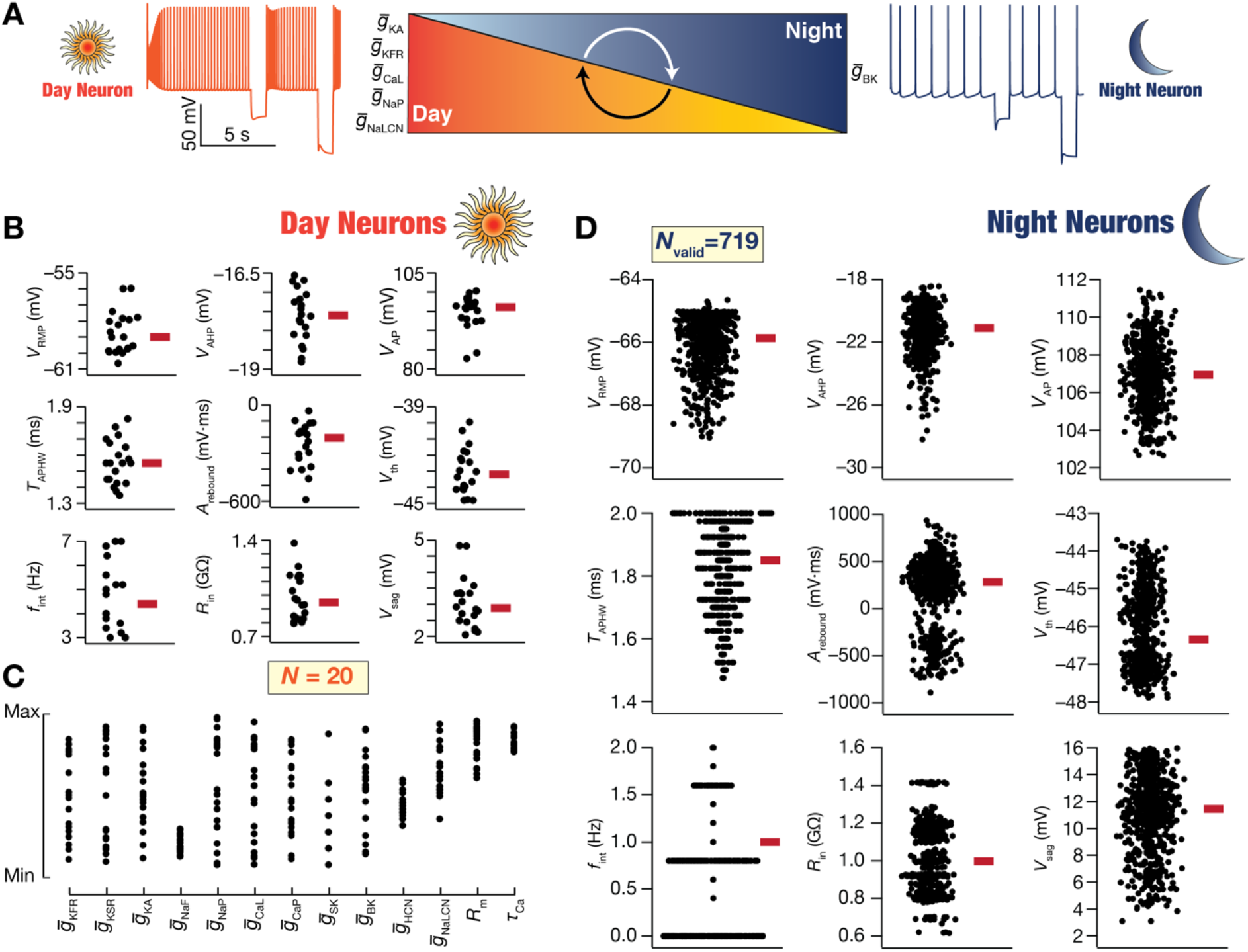
Heterogeneous distribution of measurements from night-like SCN neurons derived through unbiased search of transitions from day-like neurons. (*A*) Schematic diagram of day-to-night transitions, along with representative voltage traces from day-like and night-like models of SCN neurons. Electrophysiologically observed circadian transitions in 6 ion-channel conductances are shown. 5 conductances (*g*_Na*LCN*_, *g*_N*aP*_, *g_CaL_*, *g*_KFR_ and *g*_KA_) show high values during daytime, whereas *g*_BK_ manifests low values during daytime. (*B–C*) Beeswarm plots of measurements (*B*) and parameters (*C*) of the 20 day-like SCN neuron models that were subjected to the day-to-night transition. The transition was implemented through a modified MPMOSS algorithm that was employed to perform an unbiased search on the plasticity space. The plasticity space accounted for the physiological direction of changes in the six ion channels (shown in panel A) that are known to undergo plasticity during circadian oscillations. The widespread distribution of the measurements and the parameters of the 20 day-like neurons may be noted. (*D*) Bee-swarm plots of the measurements from 719 different SCN night-like neurons derived from the 20 day-like neurons. Red bars represent the median values. The validation process for obtaining night-like neurons employed established electrophysiological bounds on each measurement (Table 2).

Although the direction of changes in each channel subtype is known, the question of whether such transitions require *unique* magnitude of changes in these individual channels is unanswered. In addition, given the observed heterogeneities in SCN neurons recorded during day (or night) periods, how do different day (or night) neurons endowed with disparate ion channel composition achieve transitions towards achieving night-like (or day-like) characteristics? To address these questions, we picked 20 day-like models that manifested heterogeneities in the measurements (Fig. 7*B*) and underlying channel conductances (Fig. 7*C*). We first performed a stochastic search on the *plasticity space* involving changes in the subset of ion channels mentioned above for day-to-night (Fig. 1; Table 3) transitions, confined to the experimentally determined direction of change for each ion-channel subtype (Table 3; Fig. 7*A*).

For each of the 20 day-like neurons, we randomly sampled the 6-dimensional plasticity space involving sign-enforced changes in ion channel densities (Table 3). We subjected individual models to this randomized plasticity by altering the respective channel densities as per the generated sample. We measured the 9 electrophysiological properties of the models after they underwent plasticity and validated them against night-like characteristics of SCN neurons (Table 2). Plasticity parameters that yielded valid night-like neurons were declared as valid day-to-night transitions. For each of the 20 day-like neurons, we generated 200 to 1000 random samples on the plasticity space (Table 3) with a goal of generating at least 20 (max 67 from a day-like neuron) valid day-to-night transitions from each day-like neuron. This process together yielded 719 valid day-to-night transitions from a total of 8000 random transitions that were generated, with all neurons that were declared valid showing night-like characteristics (Fig. 7*D*; *cf*. Table 2). Importantly, these 719 night-like neurons manifested pronounced heterogeneities within the valid measurement ranges (Fig. 7*D*), thus providing us with a heterogeneous population of night-like neurons for further measurement as well as parametric analyses.

From the measurements perspective, as the intrinsic firing rate is lower in night-like neurons compared to their day-like counterparts (Table 2), we found that 141 of these neurons were not spontaneously active with the remaining 578 neurons showing low spontaneous firing (Fig. 7*D*; *cf*. Fig. 4*A* for day-like neurons). To maintain consistency, we measured action potential properties from spontaneous action potential firing and therefore do not report action potential properties for the silent neurons. We computed pairwise correlations between measurements across the intrinsically active (Fig. 8*A*) and silent (Fig. 8*B*) neurons separately, and found mostly weak correlations between these measurements. As mentioned earlier, the few measurements that showed strong correlations were largely owing to the relationships between the way these measurements were obtained.

**Figure 8.**
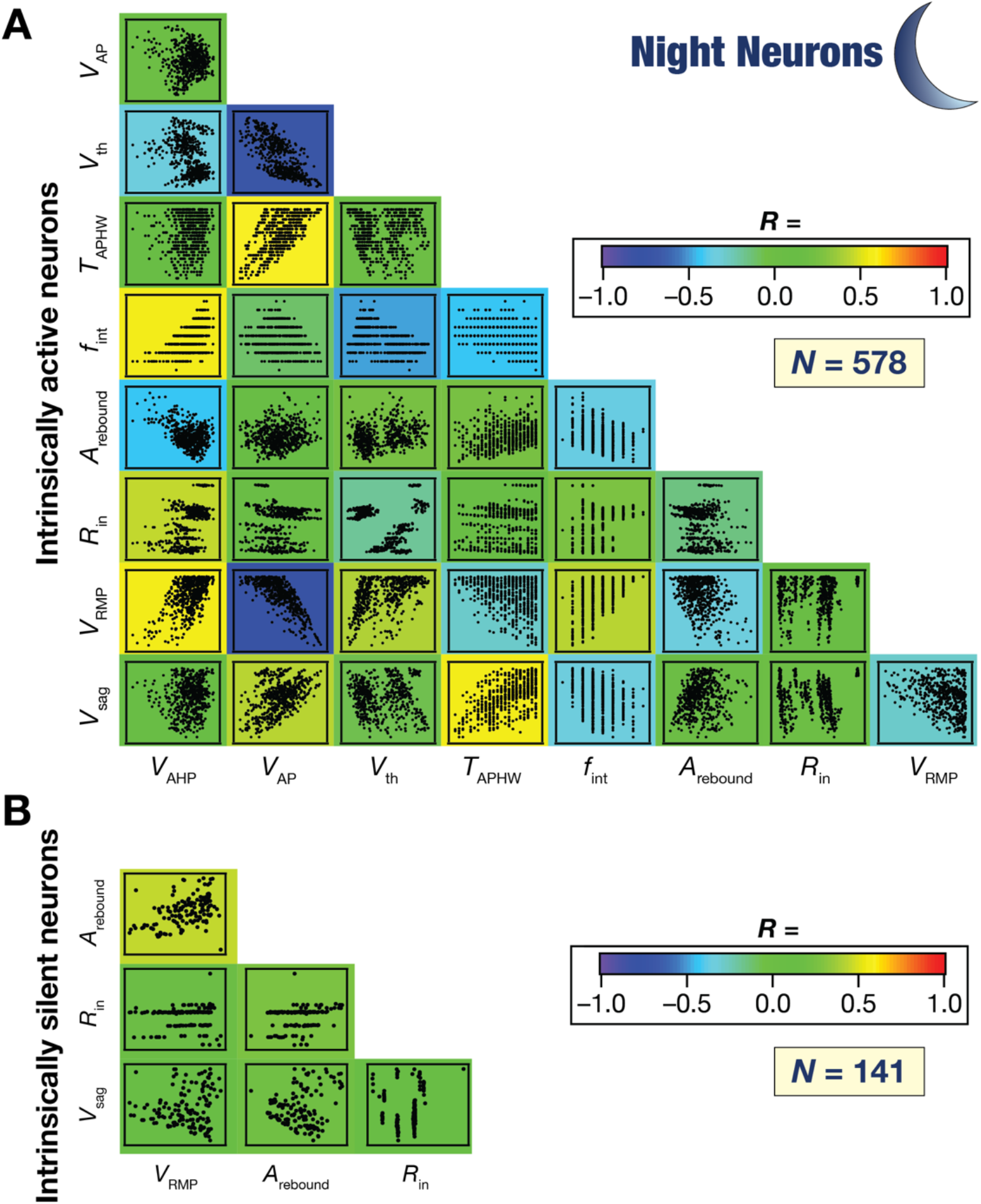
A majority of pairwise correlations between measurements from the night-like neuronal model population were weak. (*A–B*) Pair-wise correlations between the different measurements used to characterize and validate night-like neuronal models. The scatter plots are overlaid on top of the color-coded scatter plot matrix showing the respective Pearson’s correlation coefficient values. Panels *A* and *B* show measurement correlations from intrinsically active (*A*) and intrinsically silent (*B*) night neurons. The matrix in *B* does not include action potential measurements because these neurons are intrinsically silent, and action potential properties were derived from spontaneous action potentials.

The parameters associated with the ion-channel subtypes that were allowed to change during day-to-night transitions manifested widespread heterogeneities, even expanding the ranges allocated (Table 1) for the original day-like model neurons (Fig. 9*A–F*). Notably, although these 719 night-like neurons were derived from merely 20 day-like neurons by allowing only 6 of the 13 original parameters to change during the transition, there were no strong pairwise correlations between the different parameters that governed these model neurons (Fig. 9*G*). Finally, these models also showed large distances among them with both Mahalanobis (Fig. 9*H*) and normalized Euclidean (Fig. 9*I*) distance metrics. The results of linear dimensionality reduction analysis on the measurement and parametric spaces of these 719 neurons, using PCA are shown in Fig. S2. The widespread parametric distributions or the lack of pairwise relationships between model parameters should not be misconstrued that any arbitrary value of these conductances or any magnitude of sign-enforced plasticity associated with these 6 channels would yield valid day-to-night transitions or valid night models. It should be noted that most transitions (7281 of 8000=91%) were rejected in the stochastic search process that spanned this very plasticity space, and only specific *combinations* of plasticity yielded valid night models. Thus, the emphasis should be on the global structure in the parametric space whereby specific combinations of disparate plasticity profiles yielded neurons with similar functional profiles. Together, these analyses unveiled two important conclusions: (i) **ion-channel degeneracy**: disparate ion channel combinations, with no strong pairwise relationships in channel expression profiles, can yield similar characteristic night-like neurons; and (ii) **plasticity degeneracy**: combinations of very different magnitudes of sign-enforced changes involving 6 disparate ion channels can yield valid and functionally equivalent day-to-night transitions.

**Figure 9.**
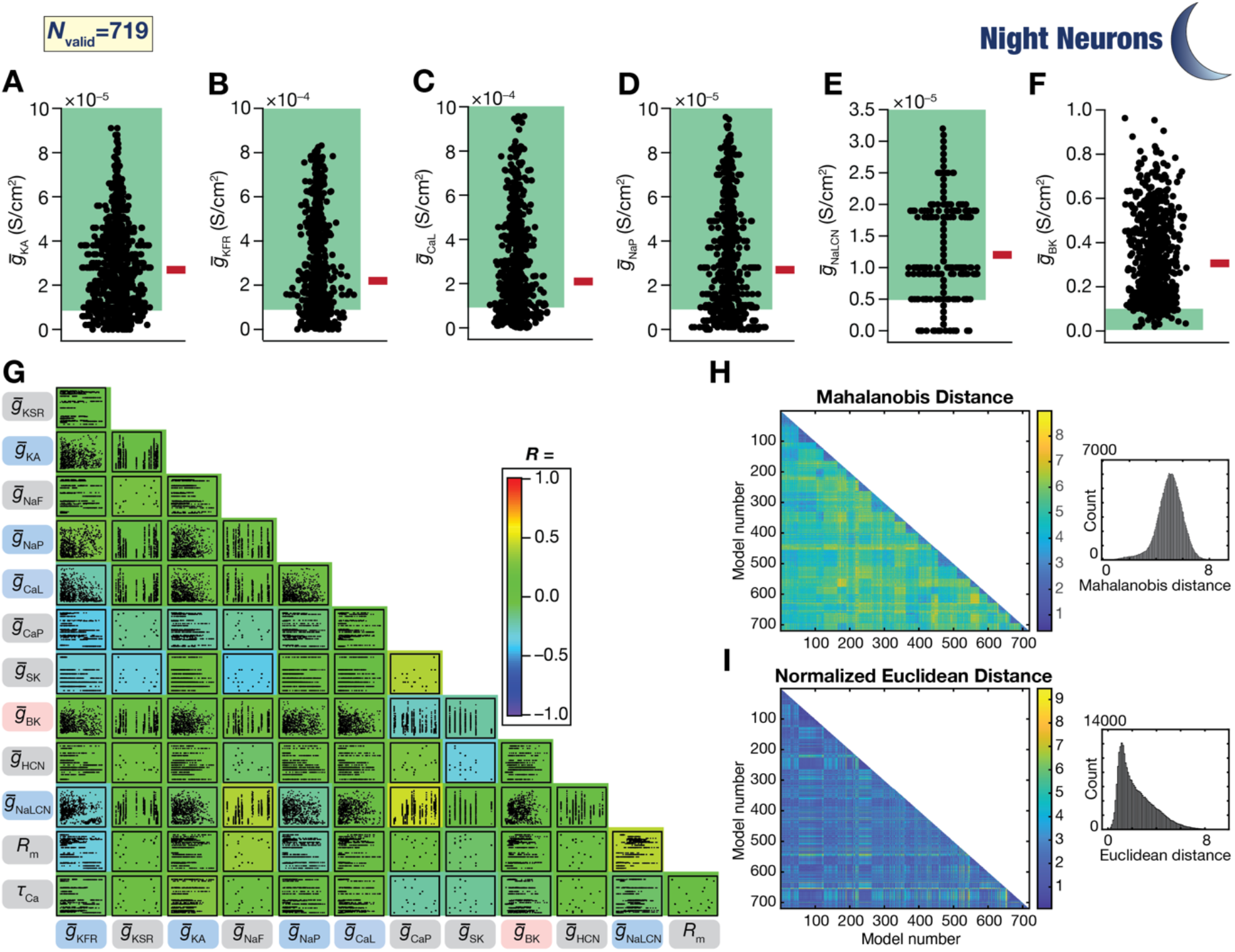
Heterogeneous distribution and weak pairwise correlations of parameters from night-like SCN neurons derived from day-like neurons. (*A–F*) Bee-swarm plots of the parameters from 719 different SCN night-like neurons shown in Fig. 7. Red bars represent the median values. Green shaded portions represent the set range of the respective parameters in the original day-neuron population (Table 1). (*G*) Scatter-plot matrix between the different parametric values of the 719 night-like SCN models shown in Fig. 7. The background color represents the value of the Pearson correlation coefficient between the different parameters. Ion channels represented with blue and red backgrounds show reductions and increases during day-to-night transitions, respectively. Parameters with gray background do not undergo any change during day-to-night transitions, and thus do not change from their respective day neurons. (*H–I*) Heterogeneities in model parameters quantified with Mahalanobis (*H*) or normalized Euclidean (*I*) distances. The matrices represent the pairwise distance between the parametric vectors defining the 719 models. Insets show the histogram of the values in the distance matrix.

### Ion-channel plasticity associated with day-to-night transitions were constrained by a low-dimensional plasticity manifold

How constrained was the measurement space, the parametric space and the fold changes associated with the valid night-like neurons and the transitions that resulted in these neurons? An ideal way to visualize constraints of a given set of data points in an *N*-dimensional space is to use dimensionality reduction techniques. We subjected nonlinear dimensionality reduction techniques on the 9-dimensional measurement space (Fig. 10*A*) and the 13-dimensional parametric space (Fig. 10*B*) associated with the 719 valid night-like neurons. We also assessed pairwise relationships between plasticity in the ion channel conductances (Fig. 10*C*) as well as a dimensionality reduction analysis on the 6-dimensional plasticity space (Fig. 10*D*) that yielded these valid night-like neurons. The night neurons were color-coded based on the specific 20 day-like neurons from where these neurons transitioned to manifest night-like characteristics (Fig. 10*A–B*, Fig. 10*D*).

**Figure 10.**
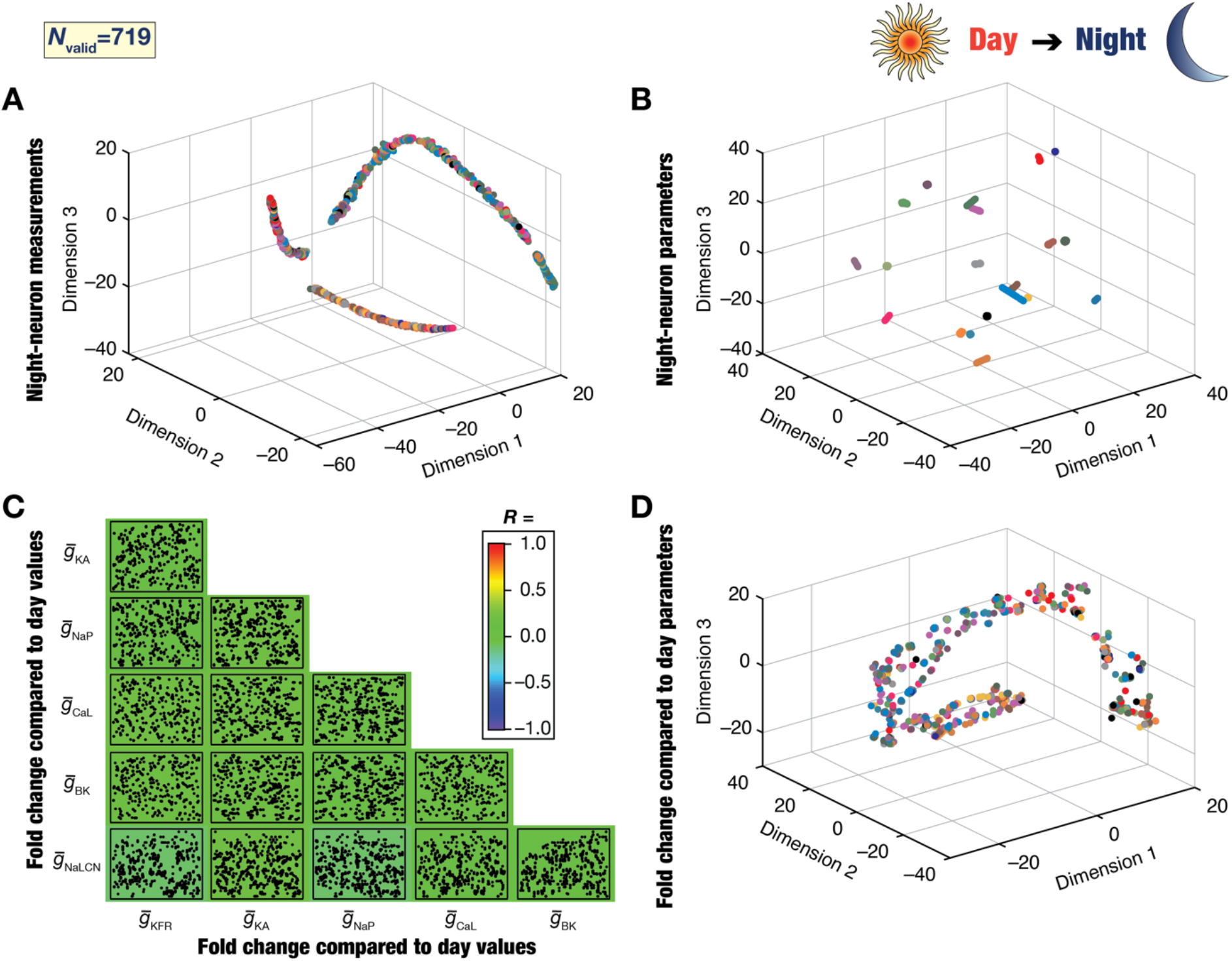
Analyses involving dimensionality reduction of fold-changes in ion-channel conductances involved in day-to-night transitions revealed the existence of plasticity manifolds. (*A*) Representation of the 9 measurements from all 719 night-like neurons on a reduced 3-dimensional space obtained through *t*-SNE. Different colors represent the 20 distinct day-like neurons from which the night-like models were obtained (shown in Fig. 7*B*). The absence of clustering based on the day-like neuron implies that night-like neurons with distinct origin may show similar measurement phenotypes. (*B*) Representation of the 13 parameters from all 719 nightlike neurons on a reduced 3-dimensional space obtained through *t*-SNE. Different colors represent the 20 distinct day-like neurons from which the night-like models were obtained (shown in Fig. 7*C*). The parameters associated with the night-like neurons formed distinct clusters based on day-like neuron where they transitioned from. (*C*) Scatter-plot matrix showing pairwise relationships between the fold changes in the six ion-channel conductances that underwent plasticity to yield the 719 valid night-like neurons from the 20 day-like neurons. The background color represents the value of the Pearson correlation coefficient between the different fold changes and indicate weak pairwise correlations across all pairs. (*D*) Representation of the fold changes in the 6 ion-channel conductances that yielded the 719 night-like neurons, with reference to conductance values in their respective day-like counterparts. Fold changes are shown on a reduced 3-dimensional space obtained through *t*-SNE. Different colors represent the 20 distinct day-like neurons from which the night-like models were obtained (ion channel distributions in day-like neurons shown in Fig. 7*C*). Note the absence of clustering based on day-neuron colors. The values spanned a small manifold within the allowed space of all transitions, illustrating a structured regime involving plasticity in different ion channels that governed day-to-night transitions.

These analyses provided several critical insights about the circadian oscillations observed in intrinsic properties of SCN neurons. First, these analyses showed that the night-like neurons occupied a low-dimensional manifold in the 9-dimensional measurement space (Fig. 10*A*). These observations demonstrated that the measurement space associated with these neurons, required to match the characteristic night-like properties of SCN neurons, was indeed constrained and did not involve arbitrary measurement combinations. Importantly, the measurement space did not manifest clusters that were defined by the specific day-neuron origin of the night-like neurons (Fig. 10*A*). This implied that the measurement space was not constrained by the specific day-like neuron where the night-like neuron originated, and manifested intermingled distribution in the reduced dimensionality space (Fig. 10*A*). Together, the measurements from night-like neurons were not distinguishable based on the origin day-like neuron and were constrained within a low-dimensional manifold in the measurement space.

Second, turning to the parametric space, we found 20 distinct clusters of parameters governing the 719 valid night-like models, with each cluster related to the 20 distinct day-like neurons where they transitioned from (Fig. 10*B*). This was in striking contrast to the similarity of the measurements observed across these neurons (Fig. 10*A*), and together provided a clear visualization of degeneracy, by showing the disparate parametric combinations that could result in the emergence of similar night-like characteristic functions (Fig. 10*A–B*). It is important to emphasize that despite the clusters observed, not all perturbations (plasticity in the 6-dimensional space) from the original day-like model were valid night-like models as a majority were declared invalid.

Finally, we asked whether the plasticity observed was constrained or occupied the entire possible range of the 6-dimensional plasticity space (in Table 3). There were no strong pairwise relationships between plasticity in any pair of channel conductances (Fig. 10*C*), suggesting the absence of correlated plasticity in conductances that resulted in valid night-like models. We performed nonlinear dimensionality reduction analysis on this 6-dimensional plasticity space, to visualize the manifestation of manifolds or clusters in plasticity space (Fig. 10*D*). Strikingly, we found that the fold changes from day-like parameters that resulted in the valid night-like parameters were constrained within a low-dimensional manifold within the plasticity space (Fig. 10*D*). This is surprising because this plasticity manifold was observed despite the heterogeneities in the origin day-like model measurements and parameters (Fig. 7), the heterogeneities in the valid night-like model measurements (Fig. 8) and parameters (Fig. 9), the presence of clusters in the night-like parametric space (Fig. 10*B*), and the absence of pairwise relationship between plasticity measurements across these models (Fig. 10*C*). Importantly, there was no clustering based on the origin day-like neuron, and valid transitions were spread throughout the plasticity manifold. Our conclusions on nonlinear dimensionality reduction analyses on the measurement, parametric, and plasticity spaces associated with these night-like neurons were invariant to the specific dimensionality reduction technique employed for the analyses (results for UMAP and PHATE on these datasets are shown in Fig. S3).

Together, despite the absence of strong correlations between plasticity in individual conductances (Fig. 10*C*), only a subset of *combination of changes forming a low-dimensional manifold* was permitted as valid day-to-night transitions, irrespective of the origin day-like models (Fig. 10*D*).

### Absence of plasticity manifolds in the night-to-day transitions

These day-to-night transitions constitute one half of the circadian oscillatory cycle. To complete the circadian cycle, we repeated the process of picking heterogeneous population of models, subjecting them to randomized sign-enforced plasticity, and validating them against electrophysiological measurements (Fig. 1). Specifically, we picked 26 night-like neurons which manifested heterogeneities in their measurements (Fig. 11*A*) and parameters (Fig. 11*B*). We picked these 26 models specifically from 5 different day-like neurons in the previous half cycle, so that we can track the transitions across the entire cycle. We subjected these 26 models to a sign-enforced stochastic search spanning a 6-dimensional plasticity space involving ionchannel conductances that are known to undergo plasticity during night-to-day transitions (Table 4). Specifically, for each of the 26 night-like neurons, we generated 1000 to 120000 random samples with a goal of generating at least 20 (max 206 from a night-like neuron) valid night-to-day transitions from each neuron. This process together yielded 1184 valid night-today transitions from a total of 273000 random transitions that were generated, with all neurons that were declared valid showing day-like characteristics (Fig. 11*C*; *cf*. Table 2). Importantly, these 1184 day-like neurons manifested pronounced heterogeneities within the valid measurement ranges (Fig. 11*C*; Fig. S4), thus providing us with a heterogeneous population of day-like neurons that have completed one full cycle of circadian oscillations.

**Figure 11.**
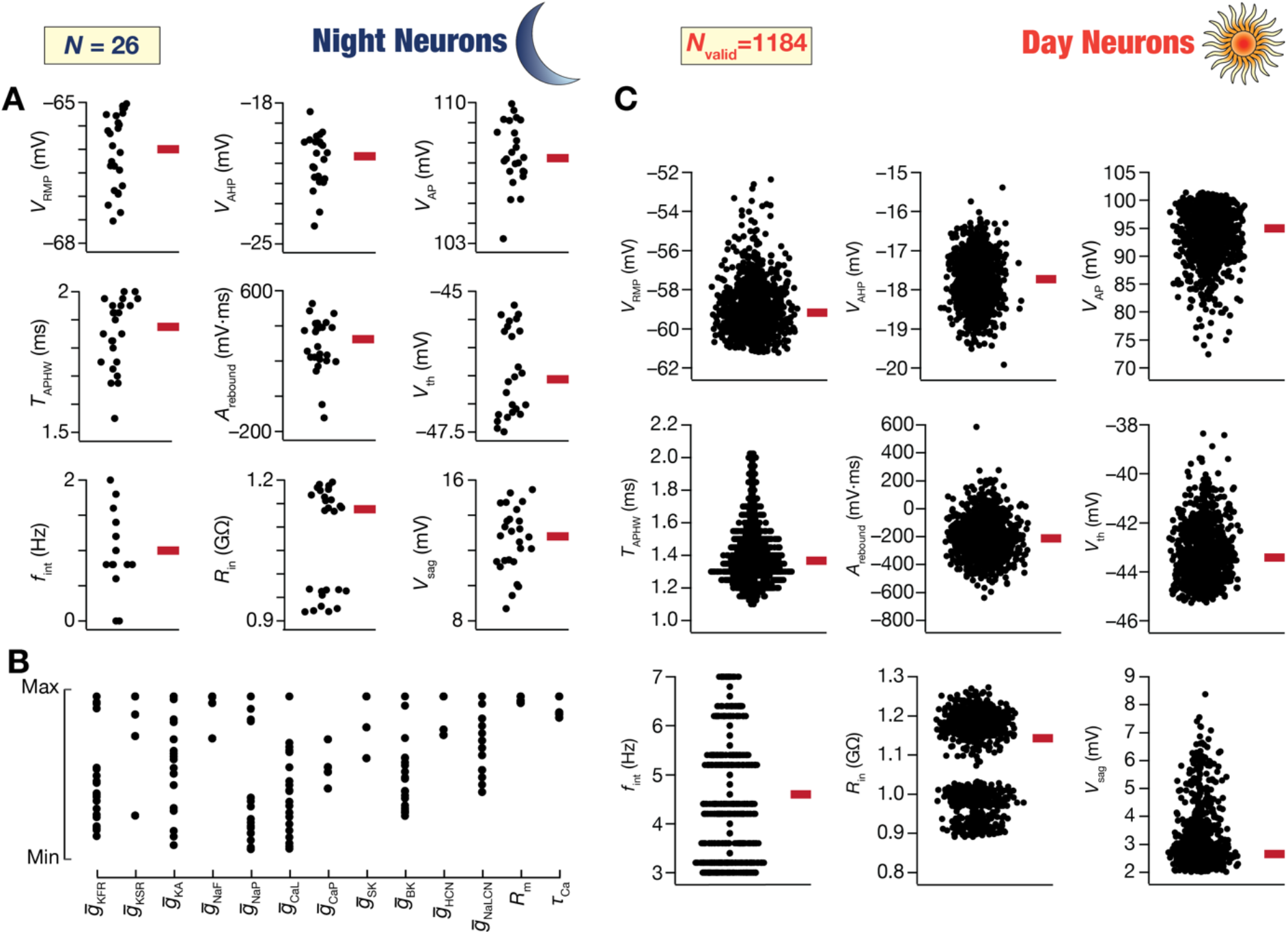
Heterogeneous distribution of measurements from day-like SCN neurons derived through unbiased search of transitions from night-like neurons. (*A–B*) Bee-swarm plots of measurements (*A*) and parameters (*B*) of the 26 night-like SCN neuron models that were subjected to the night-to-day transition. The transition was implemented through a modified MPMOSS algorithm that was employed to perform an unbiased search on the plasticity space. The plasticity space accounted for the physiological direction of changes in the six ion channels (Fig. 7*A*) that are known to undergo plasticity during circadian oscillations. The widespread distribution of the measurements and the parameters of the 26 night-like neurons may be noted. (*C*) Bee-swarm plots of the measurements from 1184 different SCN day-like neurons derived from the 26 night-like neurons. Red bars represent the median values. The validation process for obtaining day-like neurons employed established electrophysiological bounds on each measurement (Table 1).

Pairwise measurement correlations (Fig. 12) were comparable to those obtained with the original day-like measurements (Fig. 7*B*) with input resistance showing distinct clusters owing to disparate origins (Fig. 11*C*, Fig. 12). The parameter ranges for all the 6 ion-channel conductances undergoing plasticity spanned beyond the ranges specified for the generation of the initial set of valid day-like models (Fig. 13*A–F*, Fig. S5, *cf*. Table 1). The parameters that underwent plasticity showed weak pairwise correlations (Fig. 13*G*), with large distances between model parameters in the 13-dimensional parametric space (Fig. 13*H*). The distances between these models are relatively smaller (*cf*. Fig. 6*D*) because they all originated from 5 day-like neurons after completion of one full cycle of circadian oscillations with only a subset of 6 parameters changing during these transitions. The results of linear dimensionality reduction analysis on the measurement and parametric spaces of these 1184 neurons, using PCA are shown in Fig. S6.

**Figure 12.**
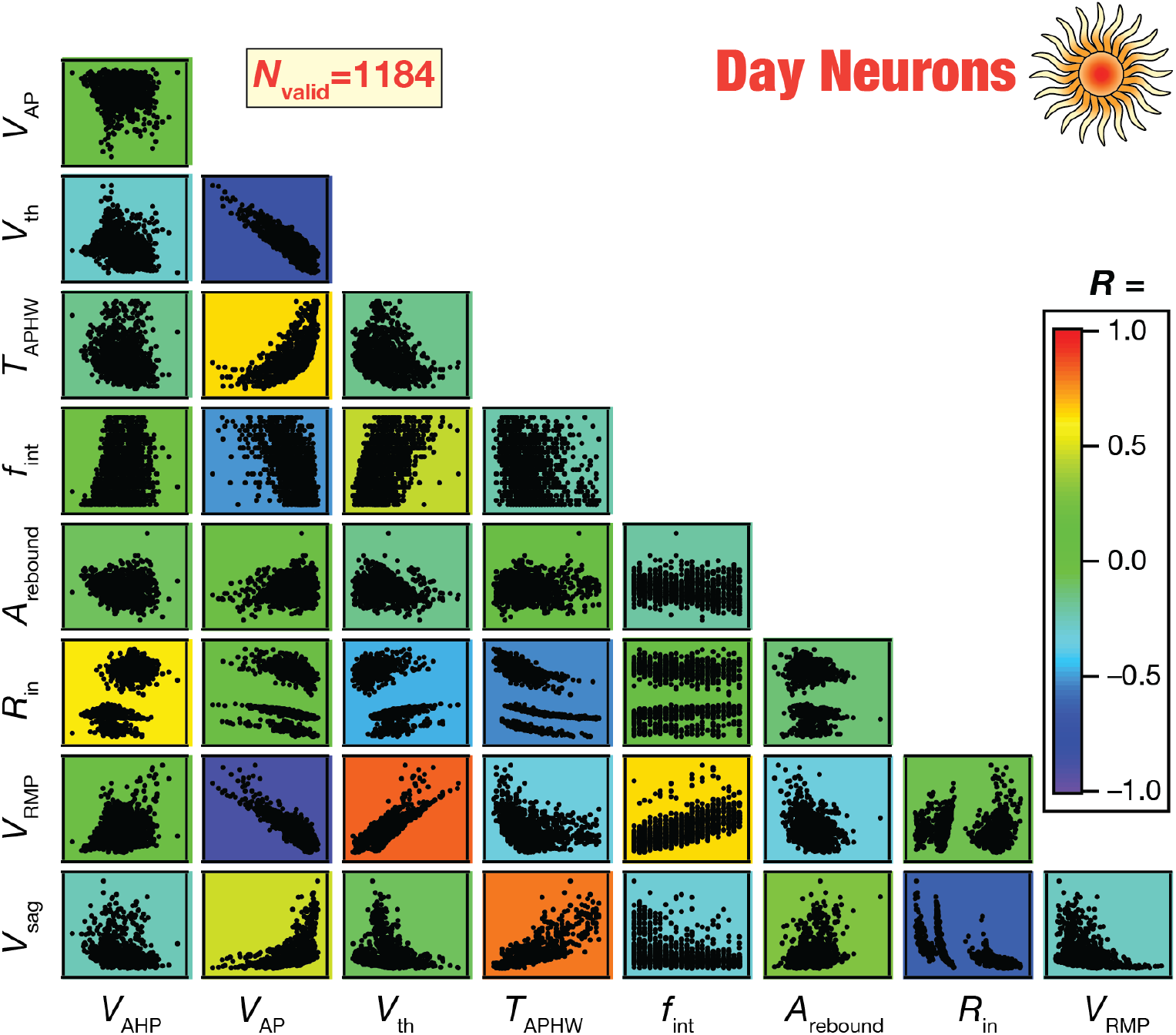
Pairwise correlations between measurements from the day-like neuronal model population derived by physiological transitions from night-like neurons. Pairwise correlations between the different measurements used to characterize and validate day-like neuronal models. The scatter plots are overlaid on top of the color-coded scatter plot matrix showing the respective Pearson’s correlation coefficient values.

**Figure 13.**
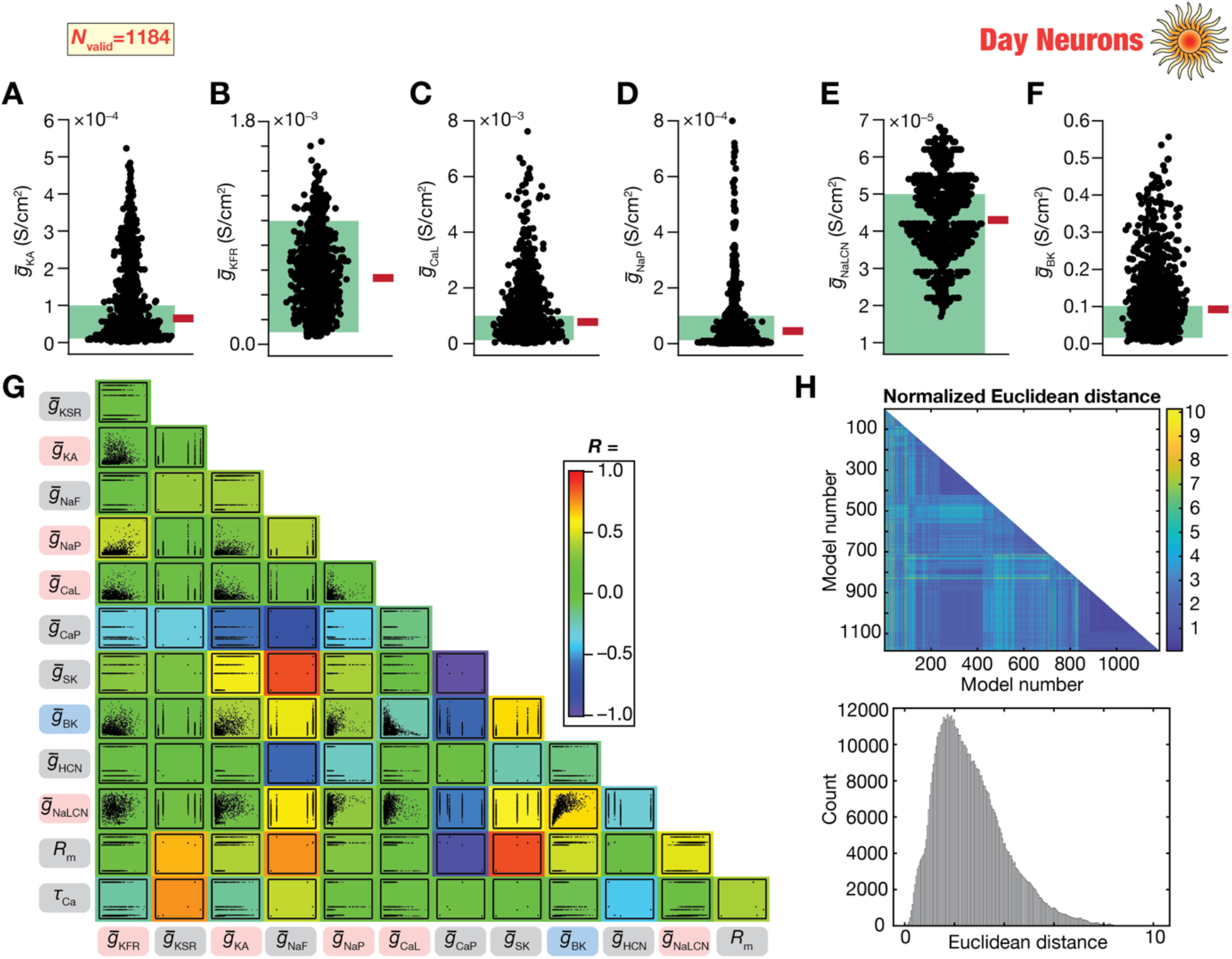
Heterogeneous distribution and weak pairwise correlations of parameters from day-like SCN neurons derived from night-like neurons. (*A–F*) Bee-swarm plots of the parameters from 1184 different SCN day-like neurons shown in Fig. 11. Red bars represent the median values. Green shaded portions represent the set range of the respective parameters in the original day-neuron population (Table 1). (*G*) Scatter-plot matrix between the different parametric values of the 1184 day-like SCN models shown in Fig. 11. The background color represents the value of the Pearson correlation coefficient between the different parameters. Ion channels represented with blue and red background show reductions and increases during night-to-day transitions, respectively. Parameters with gray background did not change during day-to-night or night-to-day transitions, and thus do not change from their respective day neurons (*N*=5). (*H*) Heterogeneities in model parameters quantified with normalized Euclidean distances. The matrix represents the pairwise distance between the parametric vectors defining the 1184 models. The bottom plot shows the histogram of the values in the distance matrix. The singularity of the associated covariance matrix precluded computation of the Mahalanobis distances for the day-like neurons.

We performed nonlinear dimensionality reduction analyses on the 9-dimensional measurement space (Fig. 14*A–B*), the 13-dimensional parametric space (Fig. 14*C–D*), and the 6-dimensional plasticity space (Fig. 14*F*) to visualize the global structure of the valid day-like neurons and the associated transitions. We found measurement manifolds in day-like neurons showing that the day-like SCN measurements were also constrained within a small subset of the 9-dimensional measurement space (Fig. 14*A–B*). Importantly, day-like models that transitioned from different night-like neurons did not manifest separate clusters but were intermingled within this low-dimensional measurement manifold showing similarity of measurements irrespective of origin neurons (Fig. 14*A–B*). On the other hand, the parametric space showed distinct clusters, with each cluster specifically related to the original day-like neuron where they originated from (Fig. 14*C–D*). Thus, similar functional outcomes were obtained from disparate combinations of underlying parameters in these day-like neurons (which have undergone one full circadian oscillatory cycle) as well. We did not observe strong pair-wise correlation between the plasticity across different ion channels that underwent plasticity during the night-to-day transitions (Fig. 14*E*). In striking contrast to the plasticity manifolds observed with day-to-night transitions (Fig. 10*F*), we did not observe a lowdimensional manifold in the 6-dimensional plasticity space associated with the night-to-day transitions (Fig. 14*F*).

**Figure 14.**
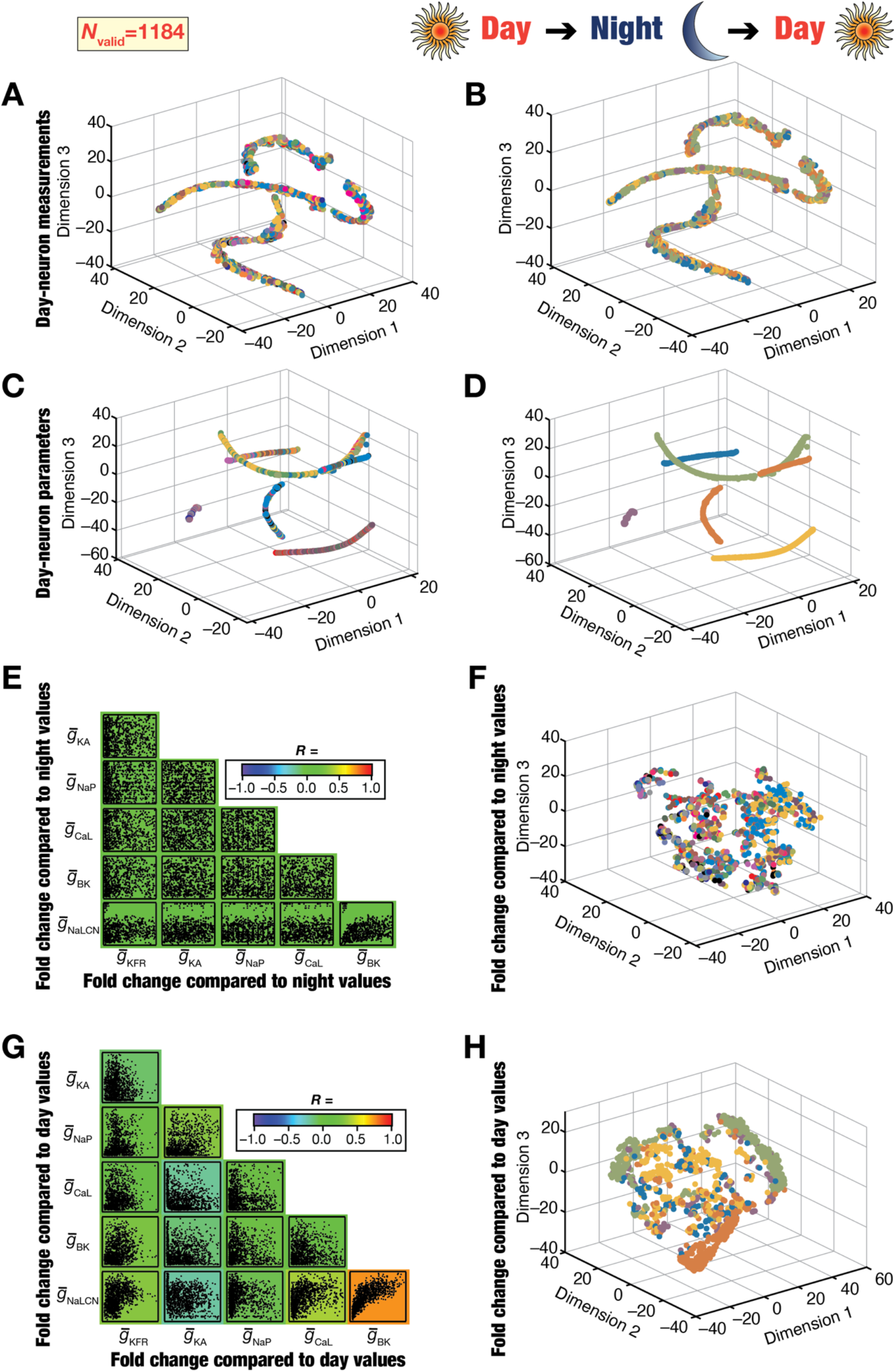
Analyses involving dimensionality reduction of fold-changes in ion-channel conductances involved in night-to-day transitions revealed the absence of structured plasticity manifolds. (*A–B*) Representation of the 9 measurements from all 1184 day-like neurons on a reduced 3-dimensional space obtained through *t*-SNE. Different colors represent the 26 distinct night-like neurons (*A*) from which the day-like models were obtained or the 5 original day-like neurons (*B*) where the 26 night-like neurons transitioned from. The absence of clustering based on the night-like neuron implies that day-like neurons with distinct origin may show similar measurement phenotypes. (*C–D*) Representation of the 13 parameters from all 1184 day-like neurons on a reduced 3-dimensional space obtained through *t*-SNE. Different colors represent the 26 distinct night-like neurons (*C*) from which the day-like models were obtained or the 5 original day-like neurons (*D*) where the 26 night-like neurons transitioned from. Day-like neurons formed distinct clusters, which were fewer than the number of distinct night neurons. (*E*) Scatter-plot matrix showing pairwise relationships between the fold changes in the six ion-channel conductances that underwent plasticity to yield the 1184 valid day-like neurons from the 26 nightlike neurons. The background color represents the value of the Pearson correlation coefficient between the different fold changes and indicate weak pairwise correlations across all pairs. (*F*) Representation of the fold changes in the 6 ion-channel conductances that yielded the 1184 day-like neurons, with reference to conductance values in their respective night-like counterparts. Fold changes are shown on a reduced 3-dimensional space obtained through *t*-SNE. Different colors represent the 26 distinct night-like neurons from which the day-like models were obtained (ion channel distributions in night-like neurons shown in Fig. 11*B*). Note the absence of clustering based on night-neuron colors. (*G–H*) Same as (*C–D*), but the fold-change analyses were performed with reference to the 5 original day-like neurons where the 26 night-like neurons were derived from, and eventually led to the 1184 day-like neurons.

As these neurons have undergone one full cycle of circadian oscillation, we assessed the cumulative plasticity undergone by these neurons through this one full cycle of oscillation. We found that there were no strong pairwise correlations between individual pairs of conductance changes (Fig. 14*G*) when fold changes in the 6 conductances of the 1184 day-neurons were compared to their 5 day-like originators. In addition, there was no plasticity manifold observed when these fold-changes were subjected to dimensionality reduction analyses, with intermingling spanning different originators across a distributed global structure (Fig. 14*H*). Our conclusions on nonlinear dimensionality reduction analyses on the measurement, parametric, and plasticity spaces associated with these day-like neurons were invariant to the specific dimensionality reduction technique employed for the analyses (results for UMAP and PHATE on these datasets are shown in Fig. S7). Our conclusions about the measurement, parametric, and plasticity spaces held even when we restricted dimensionality reduction analyses to day-like neurons that transitioned from 5 distinct night-like neurons, which in turn transitioned from 5 distinct day-like neurons (Fig. S8).

Could the manifestation of plasticity manifolds in day-to-night, but not in night-to-day, transitions be an artifact of our methodology where we first generate day neurons *de novo?* Could it be that the first transition from *de novo* neuronal population is restricted to a manifold while the second transition doesn’t manifest a manifold? To address these questions, we generated a *de novo* population of night-like SCN neurons (Fig. S9) and subjected those to night-to-day transitions to explore the manifestation of plasticity manifolds. Specifically, we generated 40,000 random models in the parametric ranges from Table 1 and subjected them to validation against night-like measurements (Table 2). We found 362 valid night-like neurons, of which we subjected 9 heterogeneous neurons (Fig. S10*A–B*) to night-to-day transitions (Fig. S9). Specifically, we explored the night-to-day plasticity space (defined by Table 4) by generating 70000 randomized transitions. We subjected each of these transitioned models to a validation process involving day-like physiological properties (Table 2) and found 290 valid day-like models from the 9 night-like neurons (Fig. S9, Fig. S10*C*). These 290 models were endowed with pronounced heterogeneities in their measurements (Fig. S10*C*) but fell within a measurement manifold in a reduced dimensional space obtained with *t*-SNE (Fig. S11*A*). The parametric space associated with these 290 day-like neurons were clustered based on the specific night-like neuron they originated (Fig. S11*B*). Importantly, there were no strong pairwise correlations between the plasticity that the 6 different ion channel densities underwent (Fig. S11*C*). There was also a striking lack of the manifestation of a plasticity manifold in the reduced dimensionality plasticity space obtained with *t*-SNE (Fig. S11*D*).

Together, these analyses (Figs. S9–S11) confirmed the absence of a plasticity manifold in night-to-day transitions, even when a *de novo* heterogeneous population of night neurons was subjected to transitions that yielded valid day-like neurons (compare Fig. S11 with *de novo* neurons with Fig. 14 obtained with transitioned neurons). These analyses suggest greater flexibility in night-to-day transitions (Fig. 14, Figs. S7–S11) compared to their day-to-night counterparts (Fig. 10, Fig. S3), with plasticity manifolds manifesting for day-to-night transitions, but not for night-to-day transitions.

## DISCUSSION

The overall experimental design involving a heterogeneous population of biophysically realistic SCN model neurons undergoing one full cycle of circadian oscillations (Fig. 1) yielded several important insights about SCN neuronal physiology. First, our analyses demonstrated the expression of ion-channel degeneracy, whereby disparate ion-channel combinations yielded SCN neurons with characteristic day- or night-like physiological properties. Second, efficacious circadian transitions in all intrinsic physiological measurements were observed despite the pronounced parametric heterogeneities that governed the day- or night-like neurons. Importantly, successful physiological transitions were feasible despite strong physiological restrictions on the identity and sign of plasticity observed in the different conductances as well as the heterogeneous expression of these conductances across different models. Third, our results unveiled plasticity degeneracy, whereby disparate combinations of sign-enforced plasticity in the 6 identified ion-channel conductances yielded valid day-to-night or night-today transitions in the intrinsic properties of SCN neurons. Plasticity degeneracy was observed even within a single neuron, where disparate combinations of parametric changes could yield valid transitions in intrinsic properties from the same origin neuron. These observations imply that at any specific phase of circadian oscillations spanning multiple cycles, individual SCN neurons should manifest pronounced parametric variability despite physiological similarity. Finally, we report the expression of plasticity manifolds, that constrain plasticity within a lowdimensional manifold within the higher dimensional plasticity space involving all possible changes, in the emergence of circadian transitions.

### Ion-channel degeneracy in the manifestation and circadian transitions of intrinsic properties: Implications for heterogeneities in SCN neuronal population

The question of how neurons in the SCN undergo circadian oscillations despite widespread heterogeneities in their biophysical composition and intrinsic properties is fundamental to understanding the precise cell-autonomous circadian changes in intrinsic properties (Yamada and Forger, 2010; Colwell, 2011; Patton and Hastings, 2018; Harvey et al., 2020). Our analyses involving a biophysically and physiologically constrained heterogeneous population of SCN neuronal models provide an elegant solution to this question from within the degeneracy framework (Edelman and Gally, 2001; Whitacre, 2010; Marder, 2011; Drion et al., 2015; Mason et al., 2015; Cropper et al., 2016; Rathour and Narayanan, 2019; Goaillard and Marder, 2021; Kamaleddin, 2022). Our analyses show that parametric heterogeneities need not be an impediment to the concomitant emergence of 9 different (Table 2) characteristic intrinsic properties of day- (Figs. 4–6, Fig. 11–13, Figs. S4–S5) or night-like (Fig. 7–9, Figs. S4–S5) SCN neurons. The ability of disparate ion-channel combinations to yield similar characteristic physiology, with these ion channels constrained by respective biophysical measurements, shows that the composition of all SCN neurons even in the same animal need not be same for them to manifest similar physiological characteristics.

In addition, although there were strong constraints on the identity of the ion channel and the direction in which they are permitted to change, our analyses show that disparate combinations of ion-channel plasticity could yield valid day-to-night (Fig. 10*C*) or night-to-day (Fig. 14*E*) transitions. These observations have several important implications. First, SCN neurons are not tightly constrained by a unique set of plasticity in the underlying channels. They are endowed with considerable freedom in choosing among several sets of transitions to make a valid physiological transition during circadian cycles. Second, if one were to follow the ion-channel composition of a single SCN neuron at a specific phase across several circadian cycles, there could be pronounced variability in the biophysical parameters across different cycles (Fig. S5) despite the physiological measurements being characteristically similar (Figs. S4). Specifically, let us say that we measure the ion-channel composition and the physiological measurements of a single SCN neuron at the same time of the day for several days. Our analyses demonstrate that while the physiological measurements across days could be identical or similar, the parametric combinations in this single neuron that yielded these physiological outcomes need not be similar. In other words, upon undergoing a full cycle, the neuron need not return to the original parametric values despite constraints on the intrinsic physiological measurements to return to similar values. We note that the possibility of pronounced parametric variability despite measurement similarity across oscillatory cycles in SCN neurons constitutes an important experimentally testable prediction within the degeneracy framework.

Third, these observations imply that the plasticity required by the same neuron to implement a valid transition (day-to-night or night-to-day) should be very different across cycles. If the parametric composition of the individual neuron changes across cycles, it stands to reason that the same magnitudes of changes that were employed in the previous cycle might not yield a valid physiological transition in this cycle. The plasticity degeneracy provides an elegant framework for achieving valid transitions not just across neurons in a population manifesting parametric heterogeneities (Anirudhan and Narayanan, 2015; Mukunda and Narayanan, 2017; Shridhar et al., 2022), but also to individual neurons transitioning across cycles through different combinations of ion-channel plasticity. Plasticity degeneracy implies that the set of ion channels and mechanisms mediating circadian oscillations could be very different in adjacent SCN neurons as well as in different cycles of the same neuron. Together, these observations emphasize the context-dependence of plasticity required to achieve valid transitions, in a manner that is tied to the current composition of the neuron undergoing the transition. Such context-dependent transitions could be further explored using computational analyses involving continual transitions as well as multi-cycle, multi-scale experimental measurements from SCN neuron cultures.

### Plasticity manifolds involving ion channels mediating circadian oscillations of intrinsic properties

Experimental lines of evidence show that SCN neurons undergo transitions from a day-like to a night-like state through concurrent changes in channel densities only in a subset of ionchannels that are expressed by them (Itri et al., 2005; Pitts et al., 2006; Itri et al., 2010; Flourakis et al., 2015; Paul et al., 2016; Whitt et al., 2018; McNally et al., 2020). It is plausible that structured rules that govern the changes in channel densities might permit only certain combinations of changes to lead to transitions. Under such a scenario, the permitted plasticity combinations would form a lower dimensional manifold in the space of allowed changes, which is termed as a plasticity manifold (Mishra and Narayanan, 2020; Mishra and Narayanan, 2021a).

We report the manifestation of plasticity manifolds in case of the day-to-night transition but not the night-to-day transition. This was visualized using nonlinear dimensionality reduction analyses, which has been widely used in the neural manifolds literature (Vyas et al., 2020), but has not been applied to analyzing manifolds in plasticity space (Mishra and Narayanan, 2021a). Our analyses demonstrate the existence of such plasticity manifolds, pointing to the presence of strong constraints on the magnitudes of sign-enforced plasticity on the ion channels. These constraints in plasticity are tightly intercoupled to several mechanisms and measurements, including the specific physiological goal(s) to be achieved, the current state of the system, the set of components that are allowed to undergo plasticity, the direction of permitted changes in the plastic component, and structure in the signaling mechanisms that govern concomitant plasticity in the components (Mishra and Narayanan, 2021a). Our analyses unveiled plasticity manifolds in day-to-night transitions despite the absence of specific signaling mechanisms that governed the concomitant changes. Our search was random and unbiased across the entire permitted plasticity space and did not impose constraints on the coupling between the plasticity in the different ion channels. Despite such independence in individual ion-channel plasticity, and despite heterogeneities in the expression of the 6 ion channels in the origin neurons, we found the manifestation of plasticity manifold in day-to-night transitions, but not in night-to-day transitions.

A simple plasticity manifold would be a scenario where plasticity of all these ion channels are co-regulated by a single mechanism, therefore resulting in pairwise correlations across all observed changes (O’Leary et al., 2013; O’Leary et al., 2014; Srikanth and Narayanan, 2015; O’Leary, 2018; Goaillard and Marder, 2021). However, our analyses show the absence of such strong pairwise correlations (Fig. 10*C*, Fig. 14*E*), instead showing the manifestation of a low-dimensional plasticity manifold (Mishra and Narayanan, 2021a) after nonlinear transformation of the plasticity space for day-to-night transitions (Fig. 10D). Experimental analyses could be employed to assess these predictions, and computational analyses involving the transcription-translation feedback loop (TTFL) could be employed to further address the structure of the plasticity manifold that results in circadian transitions.

The manifestation of a plasticity manifold in day-to-night transition but not in the nightto-day transitions emphasize the asymmetry in the mechanisms underlying the bidirectionality of physiological transitions. Although there are lines of evidence for asymmetry on the impact of bidirectional changes in other contexts (Phillips and Hasenstaub, 2016; Ebsch and Rosenbaum, 2018), our analyses suggest stringent constraints on day-to-night transitions to their night-to-day counterparts. The differential flexibilities for the two transitions emanate from the specific physiological targets (Table 2) and the permitted plasticity space involving directions of the subset of ion channels (Table 3 *vs*. Table 4). Future experimental and computational studies could examine the existence as well the mechanistic bases behind such differential constraints in the two transitions.

### Limitations and future directions

Our analyses unveiled important insights about how a heterogeneous population of SCN neurons could manifest characteristic properties and undergo signature physiological transitions across a circadian cycle. However, there are important model considerations that need to be addressed in future studies. First, our analyses are based on a simple, single-compartmental conductance-based model of an SCN neuron. Future studies could look to incorporate additional structural heterogeneities, as well as heterogeneities in dendritic ion channel distributions, in a morphologically realistic model population that manifests characteristic somato-dendritic properties. Such computational analyses have to be accompanied by morphological and electrophysiological measurements of SCN neurons across the circadian cycle. Models constrained by these experimental measurements could be employed to probe questions about the impact of structural and biophysical heterogeneities on characteristic physiological properties and their signature circadian transitions.

Second, circadian oscillations in the SCN are studied across multiple scales: the cell-autonomous SCN, the molecular mechanisms and signaling networks underlying continual circadian transitions involving the transcription-translation feedback loop (TTFL) in SCN cells, the SCN as a cellular network and, finally, the SCN as circadian orchestrator (Yamada and Forger, 2010; Colwell, 2011; Schmutz et al., 2014; Flourakis et al., 2015; Kudo et al., 2015; Paul et al., 2016; Patton and Hastings, 2018; Harvey et al., 2020). Our study focused on the ionic basis of cell-autonomous changes in neural excitability. The insights here about plasticity manifolds and ion-channel degeneracy could be employed to understand the molecular mechanisms and the role of TTFL in regulating these ion channels in the heterogeneous population of SCN cells. As one of the physiological goals is to regulate the firing rate of the neurons over the day-night cycles, these questions could be addressed from the perspective of how these goals are continually achieved while accounting for heterogeneities not just in the ion-channel composition but also in the signaling components associated with the TTFL. Importantly, a logical next step would be to assess the impact of ion-channel degeneracy and plasticity manifolds in individual neurons on synchronization of cellular oscillations throughout a heterogeneous SCN network. The converse, on how connectivity and synchronization across different neurons could affect the plasticity manifolds that govern the circadian transitions, also constitutes an important question in terms of how the network inputs define ion-channel degeneracy and plasticity manifolds in individual neurons. Such analyses would also provide insights about potential asymmetries between the day-to-night and the night-to-day transitions and about our prediction on differential expression of plasticity manifolds across these two transitions.

Third, while the cell-autonomous transitions in intrinsic properties and network synchrony through connections form one layer of complexity, an additional layer of complexity arises from the expression of several neuromodulators that influence action potential activity and rhythmicity in the SCN. These neuromodulators regulate the synchronization properties, the baseline excitability, and the plasticity of the circadian neural code with multi-scale impact of SCN physiology. In addition to accounting for heterogeneities in ion channels, calcium regulatory mechanisms, signaling cascades, neurotransmitters and their receptors, and network connectivity, SCN analyses should include heterogeneities in neuromodulatory receptor expression across SCN neurons. Future studies could investigate the impact of multi-scale heterogeneity (spanning molecular, cellular, and network scales) and explore the expression of degeneracy and plasticity manifolds across different scales while accounting for neuromodulation and light entrainment of the SCN network (Vasalou and Henson, 2010; Yamada and Forger, 2010; Colwell, 2011; DeWoskin et al., 2015; Enoki et al., 2017; Harvey et al., 2020; Mishra and Narayanan, 2021a). The SCN is an ideal system to address questions on long-term plasticity manifolds and degeneracy across scales, using both experimental and computational techniques, given the well-defined function and the tight coupling observed in the physiology and mechanisms across scales. Strong experimentally constrained multi-scale mechanistic models of the SCN can provide deeper insights into how heterogeneities, degeneracy and plasticity manifolds across different scales interact to provide precise physiology and state-dependent transitions during circadian oscillations (Townsley et al., 2020; Mishra and Narayanan, 2021a).

Finally, there are reports of changes to ion-channel properties and intrinsic properties spanning different neuronal subtypes under several pathophysiological and altered behavioral conditions (Kullmann and Waxman, 2010; Terzic and Perez-Terzic, 2010; Cannon, 2021; Deng and Klyachko, 2021; Franklin and Laubham, 2021; Gripp et al., 2021; Kessi et al., 2021; Mantegazza et al., 2021). Thus, future studies should explore the impact of altered channel properties/densities under different behavioral and pathological conditions on specific ion channels and the physiology of SCN neurons, as well as the impact on transitions and plasticity manifolds associated with them. Such studies might be performed by systematically removing ion channels from the models using virtual-knockout model analyses (Rathour and Narayanan, 2014; Anirudhan and Narayanan, 2015; Mukunda and Narayanan, 2017; Basak and Narayanan, 2018b; Mittal and Narayanan, 2018; Seenivasan and Narayanan, 2020; Mishra and Narayanan, 2021b; Roy and Narayanan, 2021) as well as by examining the electrophysiological properties of the SCN channels and neurons spanning the circadian cycle. These analyses could yield insights into the relative contribution of individual channels to circadian oscillations and unveil the impact of ion channelopathies on circadian oscillations across neurological disorders.

Together, our analyses suggest the expression of ion-channel degeneracy and plasticity manifolds as efficacious substrates for SCN neurons to undergo physiologically precise circadian transitions in their intrinsic properties despite pronounced heterogeneities in their ion-channel composition. Given the inherent heterogeneities in neuronal properties and molecular mechanisms as well as the precise long-duration cycles in characteristic measurements, the SCN constitutes an ideal system to assess ion-channel degeneracy and multi-scale plasticity manifolds spanning multiple cycles. Our analyses provide a theoretical framework that could be employed to effectively understand the intricately connected multiscale circadian oscillations in the SCN under physiological and pathological conditions.

## ACKNOWLEDGMENTS

The authors thank members of the cellular neurophysiology laboratory for helpful discussions and for comments on a draft of this manuscript. This work was supported by the DBT-Wellcome Trust India Alliance (Senior fellowship to R. N.: IA/S/16/2/502727) and the Kishore Vaigyanik Protsahan Yojana (H. N.).

## Author Contributions

H.N. and R.N. designed experiments; H.N. performed experiments; H.N. analyzed data; H.N. and R.N. wrote the paper.

## Competing Interest Statement

The authors declare that they have no competing interests.

## Notes

### Competing Interest Statement

The authors have declared no competing interest.

