## Supplementary Figures S1-S11 for "Plasticity manifolds and ion-channel degeneracy govern circadian oscillations of neuronal intrinsic properties in the suprachiasmatic nucleus"

### Supplementary Material

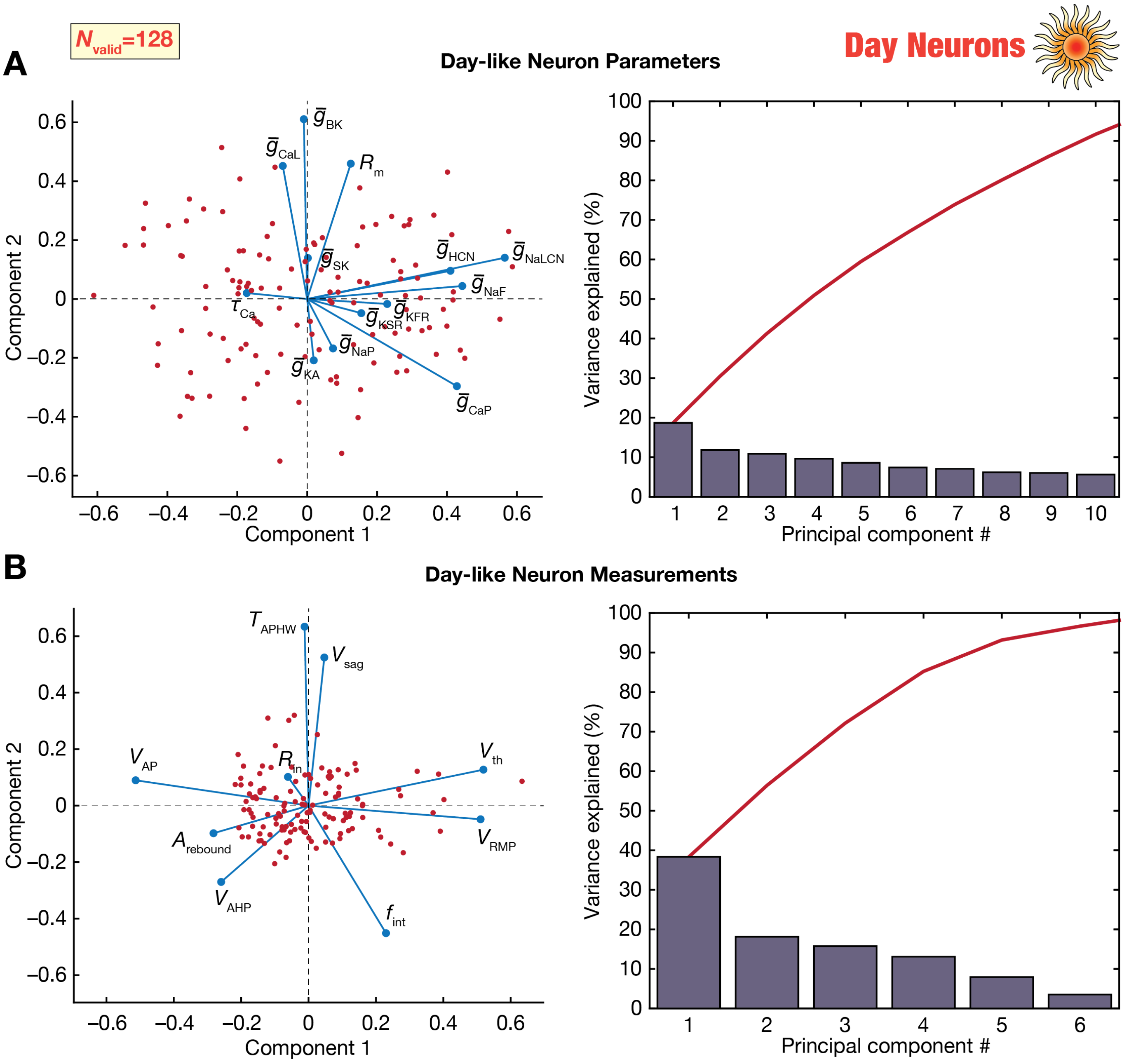

**Supplementary Figure S1**. (*A*) *Left*, Representation of the parameters of the 128 day-like neurons on a reduced 2D space obtained from PCA. The axes represent the first 2 principal components, with each point (in red) representing a distinct day-like neuron. Blue lines represent the loadings of the different parameters. *Right*, Scree plot representing the population variance explained by successive principal components. (*B*) Same as panel *A*, but for measurements of the 128 day-like neurons.

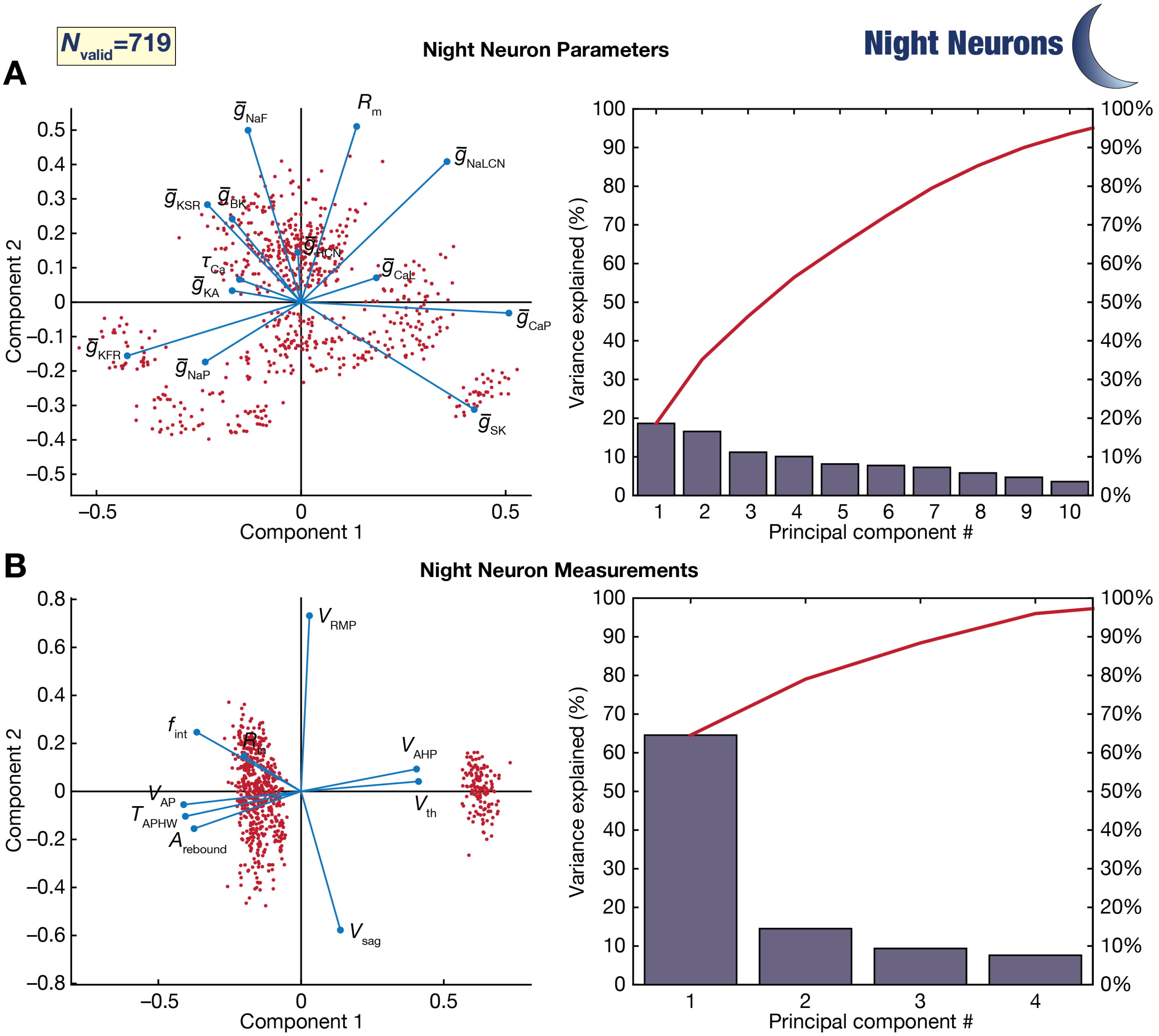

**Supplementary Figure S2**. (*A*) *Left*, Representation of the parameters of the 719 night-like neurons on a reduced 2D space obtained from PCA. The axes represent the first 2 principal components, with each point (in red) representing a distinct night-like neuron. Blue lines represent the loadings of the different parameters. *Right*, Scree plot representing the population variance explained by successive principal components. (*B*) Same as panel *A*, but for measurements of the 719 night-like neurons.

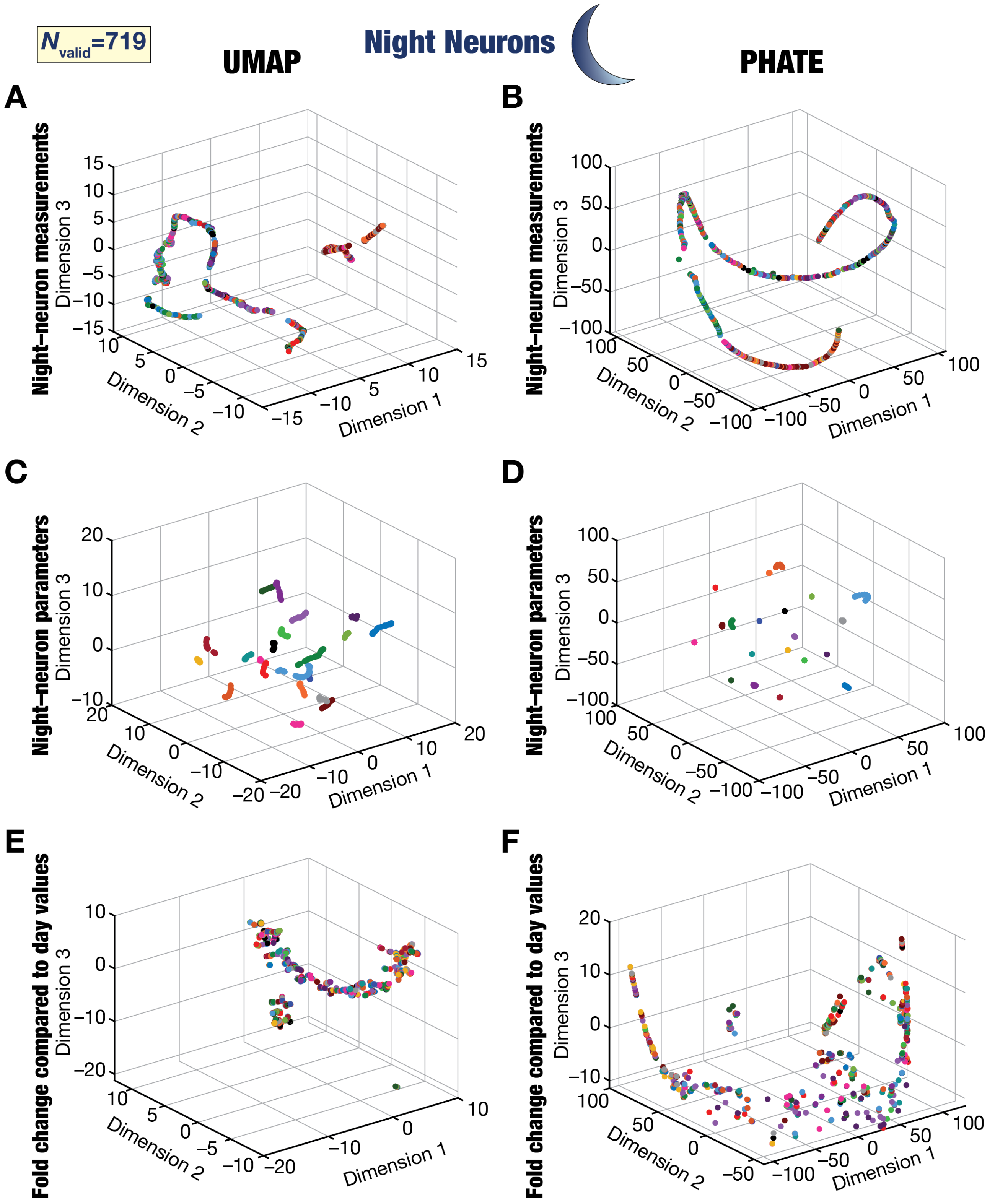

**Supplementary Figure S3.** Visualization of the measurement (*A*–*B*), parametric (*C*–*D*), and plasticity (*E*–*F*) spaces associated with the 719 night-like neuron measurements using dimensionality reduction analyses performed with UMAP (*A*, *C*, *E*) or PHATE (*B*, *D*, *F*). Each point represents a distinct night-like neuron and are color-coded (20 colors) with reference to the day-like neuron from which it transitioned.

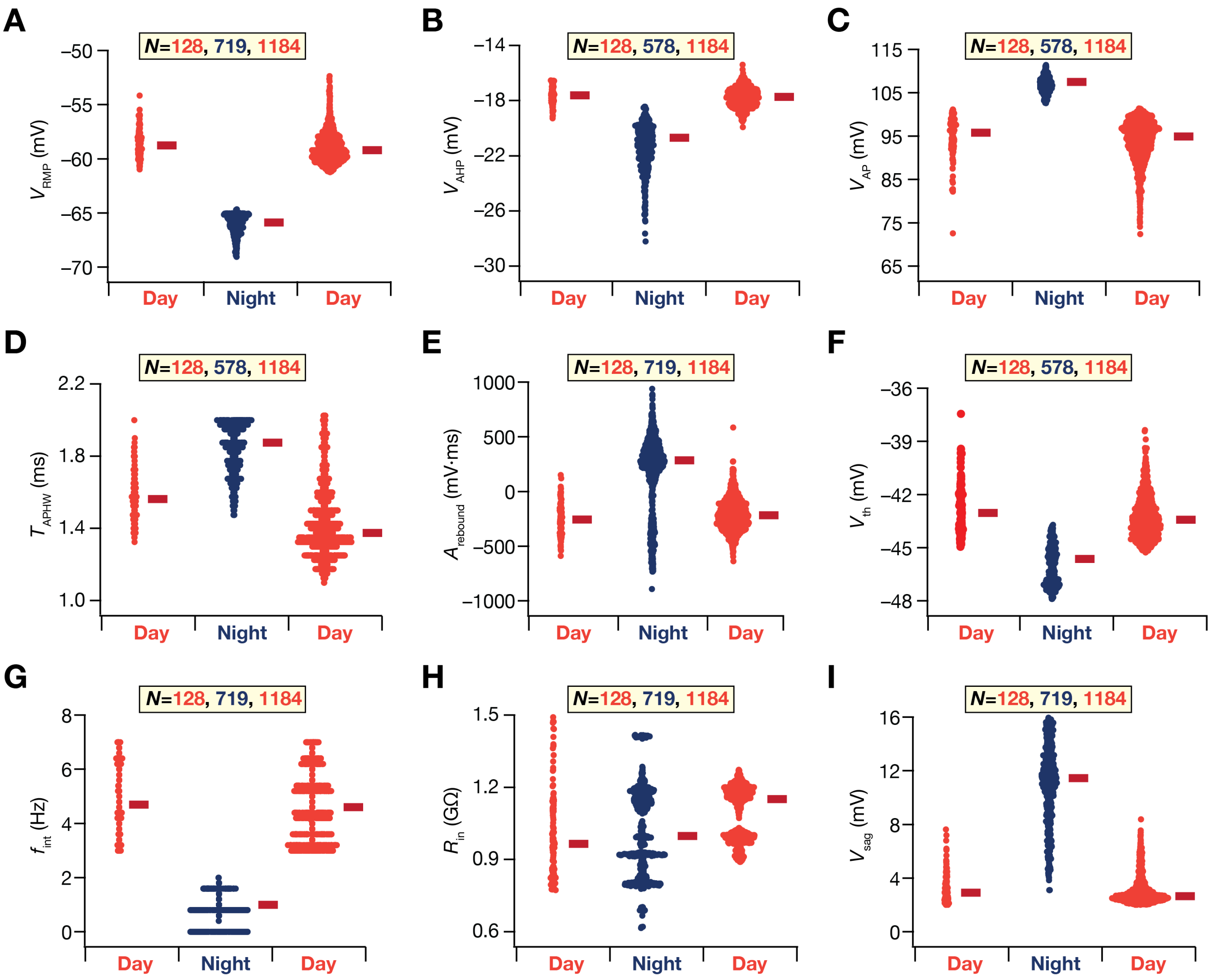

**Supplementary Figure S4**. Bee-swarm plots of all 9 physiological measurements of the different day (red) and night (blue) neurons. Figure 1 provides the flowchart of how day neurons and day-to-transition as well as night-to-day were executed within our modeling. The red line represents the median value. A clear difference in measurements from day and night neurons may be observed (*cf.* Table 2).

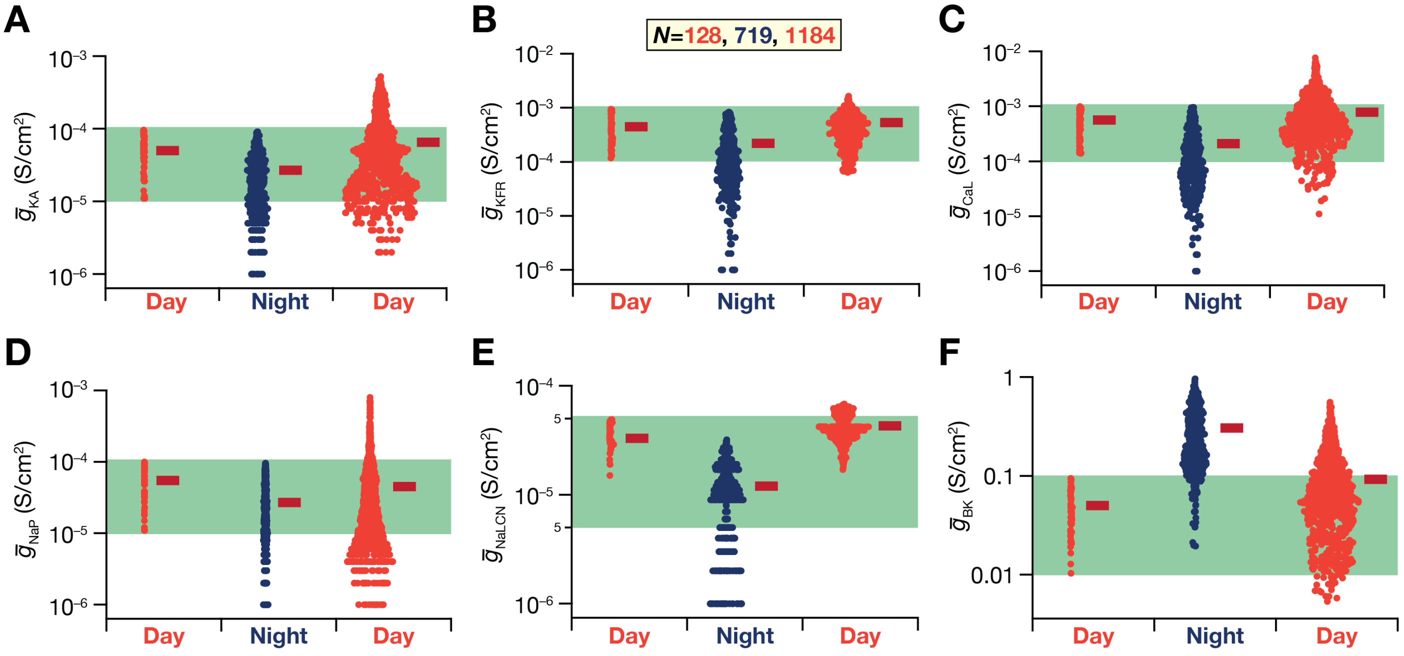

**Supplementary Figure S5**. Bee-swarm plots of the plastic parameters that were subjected to sign-enforced change during the circadian oscillations of the day (red) and night (blue) neurons. Red line represents the median. The shaded region (green) represents the bounds of the MPMOSS used to generate the initial day population (from Table 1). Figure 1 provides the flowchart of how day neurons, day-to-night as well as night-to-day transitions were executed within our modeling framework.

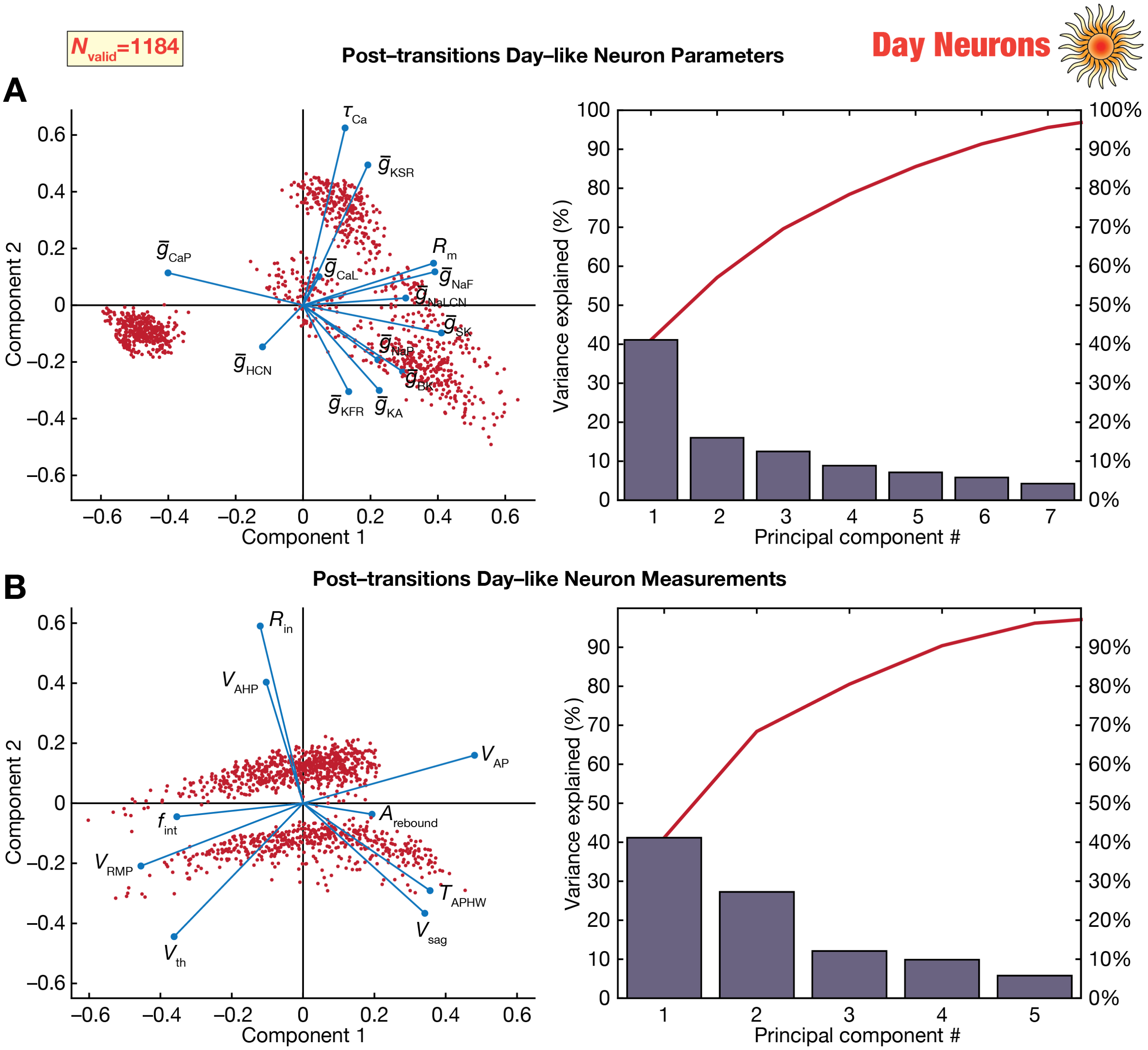

**Supplementary Figure S6**. (*A*) *Left*, Representation of the parameters of the 1184 day-like neurons on a reduced 2D space obtained from PCA. The axes represent the first 2 principal components, with each point (in red) representing a distinct day-like neuron. Blue lines represent the loadings of the different parameters. *Right*, Scree plot representing the population variance explained by successive principal components. (*B*) Same as panel *A*, but for measurements of the 1184 day-like neurons.

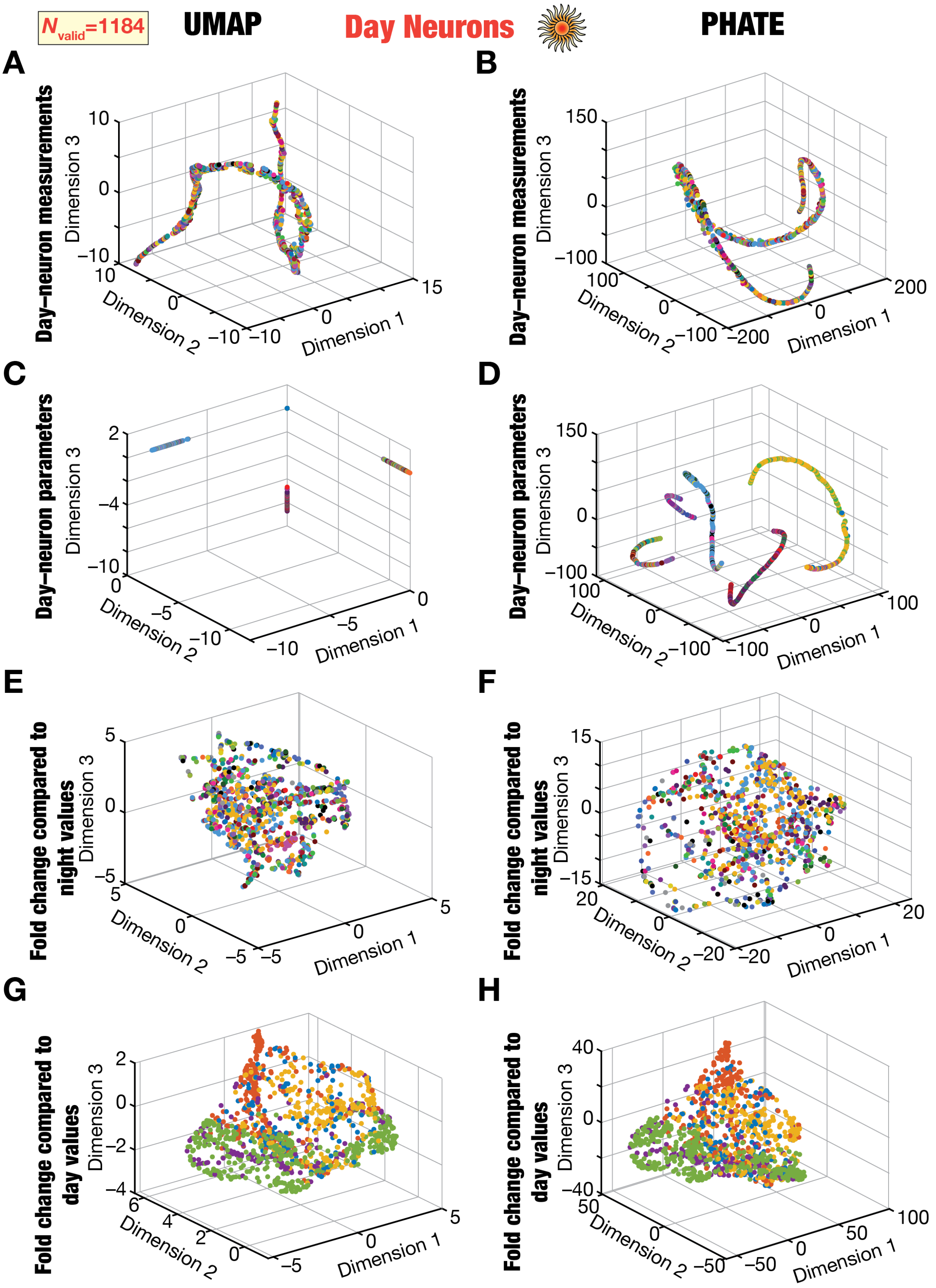

**Supplementary Figure S7.** Visualization of the measurement (*A–B*), parametric (*C–D*), and plasticity (*E–F*) spaces associated with the 1184 day-like neuron measurements (obtained after day-to-night and a subsequent night-to-day transitions) using dimensionality reduction analyses performed with UMAP (*A*, *C*, *E*, *G*) or PHATE (*B*, *D*, *F*, *H*). Each point represents a distinct day-like neuron and are color-coded with reference to the night-like neuron (*A–F*; 26 colors) or the original day-like neuron (*G–H*; 5 colors) from which it transitioned.

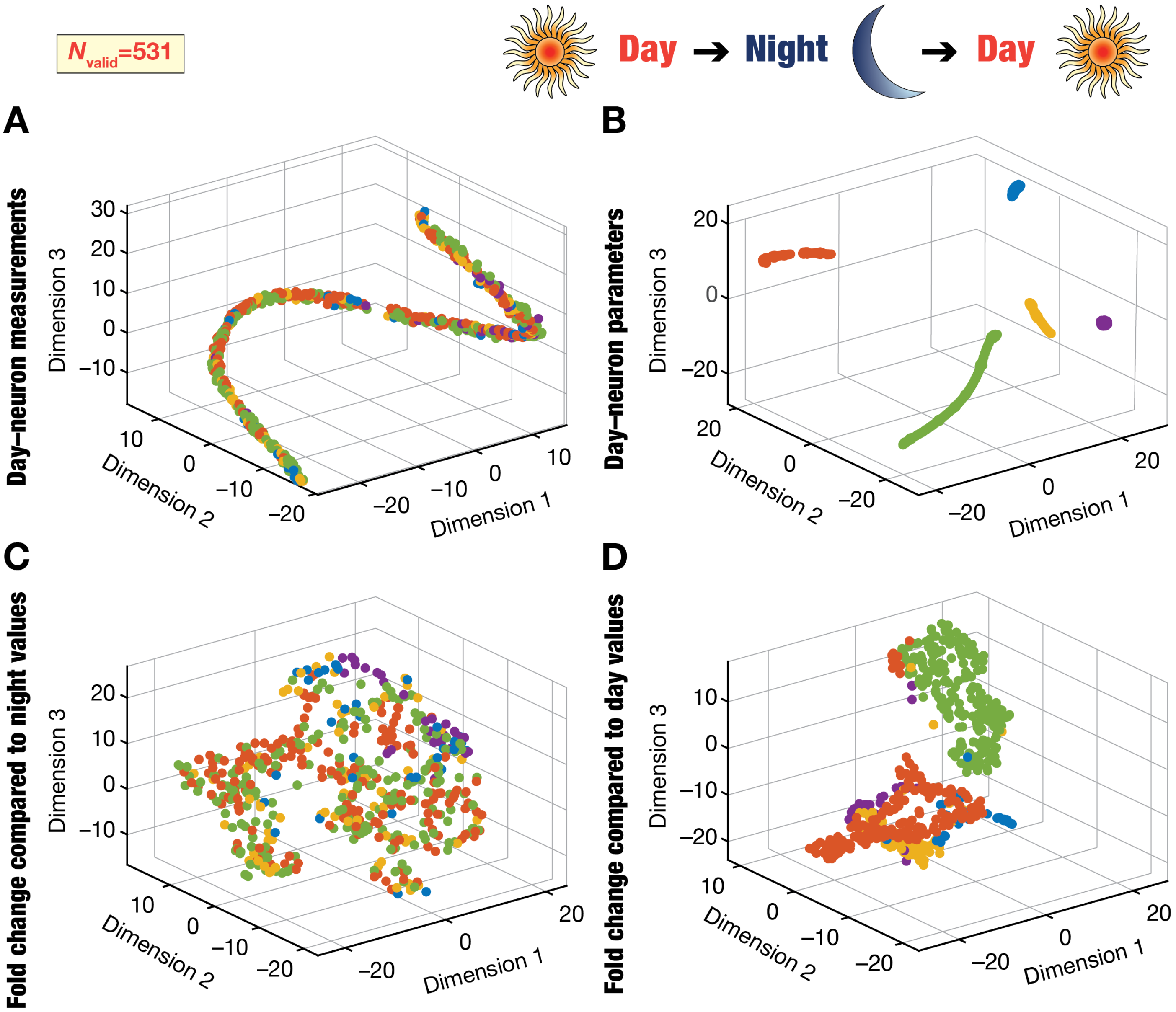

**Supplementary Figure S8.** Visualization of the measurement (*A*), parametric (*B*), and plasticity (*C*) spaces associated with 531 day-like neuron measurements (obtained after day-to-night and a subsequent night-to-day transitions) using dimensionality reduction analyses performed with *t*-SNE. These day-like neurons were obtained from 5 night-like neurons, each of which were derived from a distinct day-like neuron. The day-to-night and night-to-day transitions were implemented using the same algorithm shown in Figure 1. Each point represents a distinct day-like neuron and are color-coded with reference to the night-like neuron (*A*–*F*; 5 colors) or the original day-like neuron (*G*–*H*; 5 colors) from which it transitioned from.

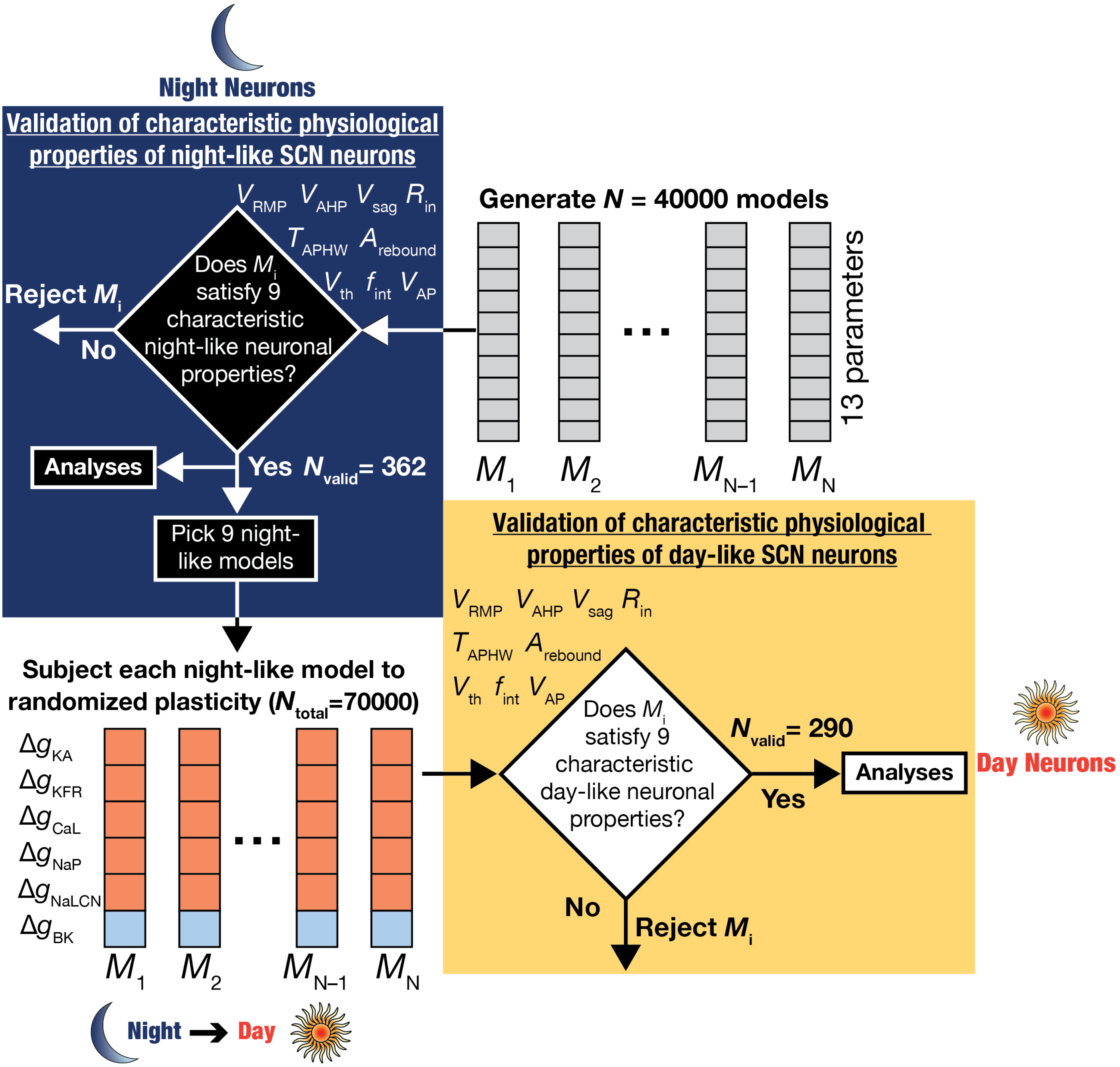

**Supplementary Figure S9.** Flowchart demonstrating the overall methodological plan for assessing ion-channel degeneracy and plasticity manifolds in circadian oscillations in SCN neurons with the cycle starting at a heterogeneous population of night-like neurons. *Left,* The first set of night-like neurons were generated by a *de novo* unbiased search involving 13 different parameters involving 40000 neuron models. Of these, 362 were found to show valid night-like physiological properties. As a second step, 9 of these 362 night-like models were picked and subjected to night-to-day transitions that involved plasticity in six different ion channels in electrophysiologically determined directions (red implies increase, blue implies reduction). *Right,* Models subjected to randomized plasticity ($N_{total}$=70000 total random transitions) were validated with day-like measurements from SCN neurons, and 290 models derived from the 9 day-like models were found to be valid. The specific combinations of ion-channel plasticity (from respective night-like neurons) that resulted in valid day-like models were subjected to dimensionality reduction analysis to determine the presence of structured plasticity manifolds in the night-to-day transitions.

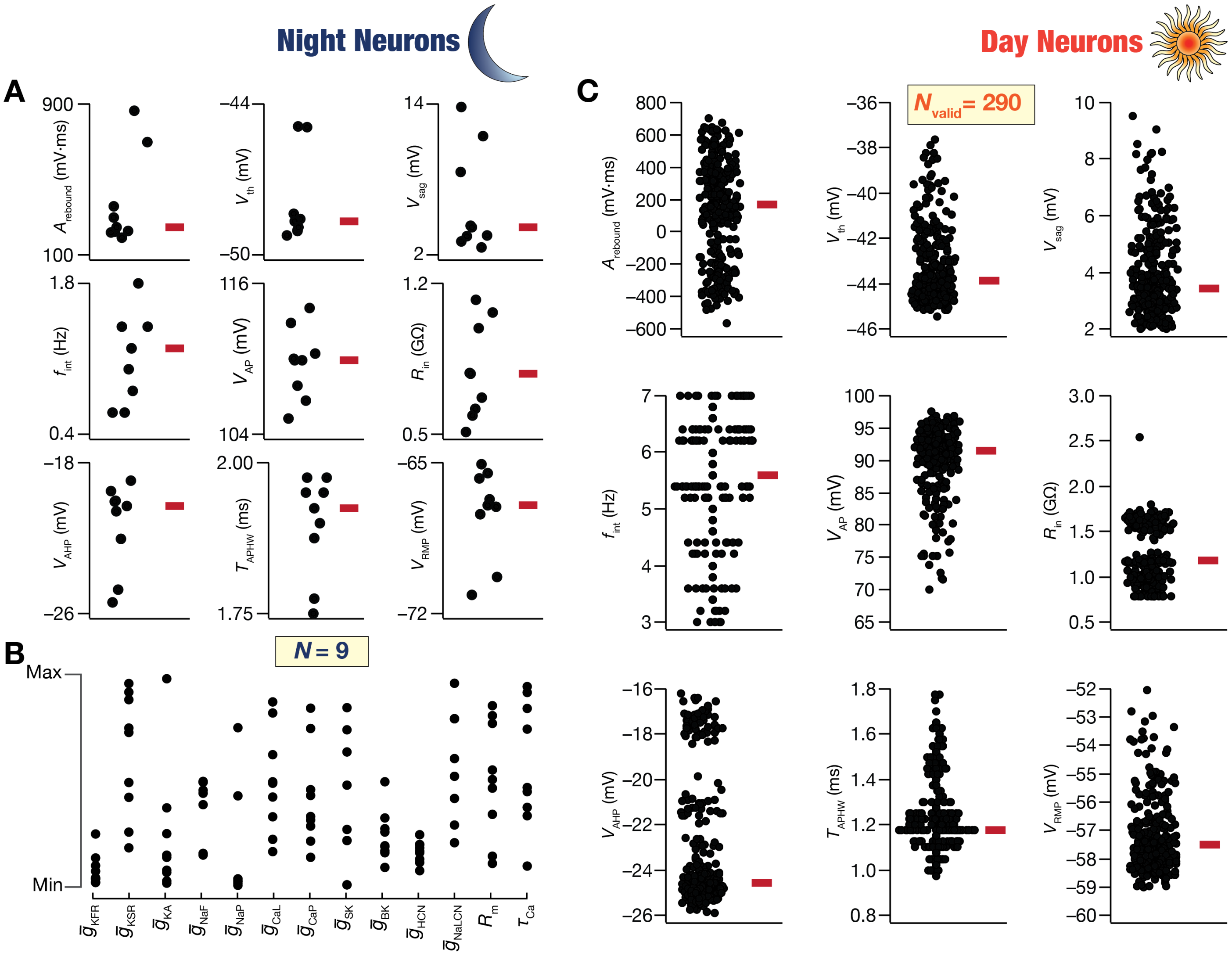

**Supplementary Figure S10. Heterogeneous distribution of measurements from day-like SCN neurons derived through unbiased search of transitions from the *de novo* population of night-like neurons.** (*A–B*) Bee-swarm plots of measurements (*A*) and parameters (*B*) of the 9 night-like SCN neuron models that were subjected to the night-to-day transition (see Supplementary Fig. S9). The transition was implemented through a modified MPMOSS algorithm that was employed to perform an unbiased search on the plasticity space. The plasticity space accounted for the physiological direction of changes in the six ion channels (shown in Supplementary Fig. S9) that are known to undergo plasticity during circadian oscillations. The widespread distribution of the measurements and the parameters of the 9 night-like neurons may be noted. (*C*) Bee-swarm plots of the measurements from 290 different SCN day-like neurons derived from the 9 night-like neurons. Red bars represent the median values. The validation process for obtaining day-like neurons employed established electrophysiological bounds on each measurement (Table 2).

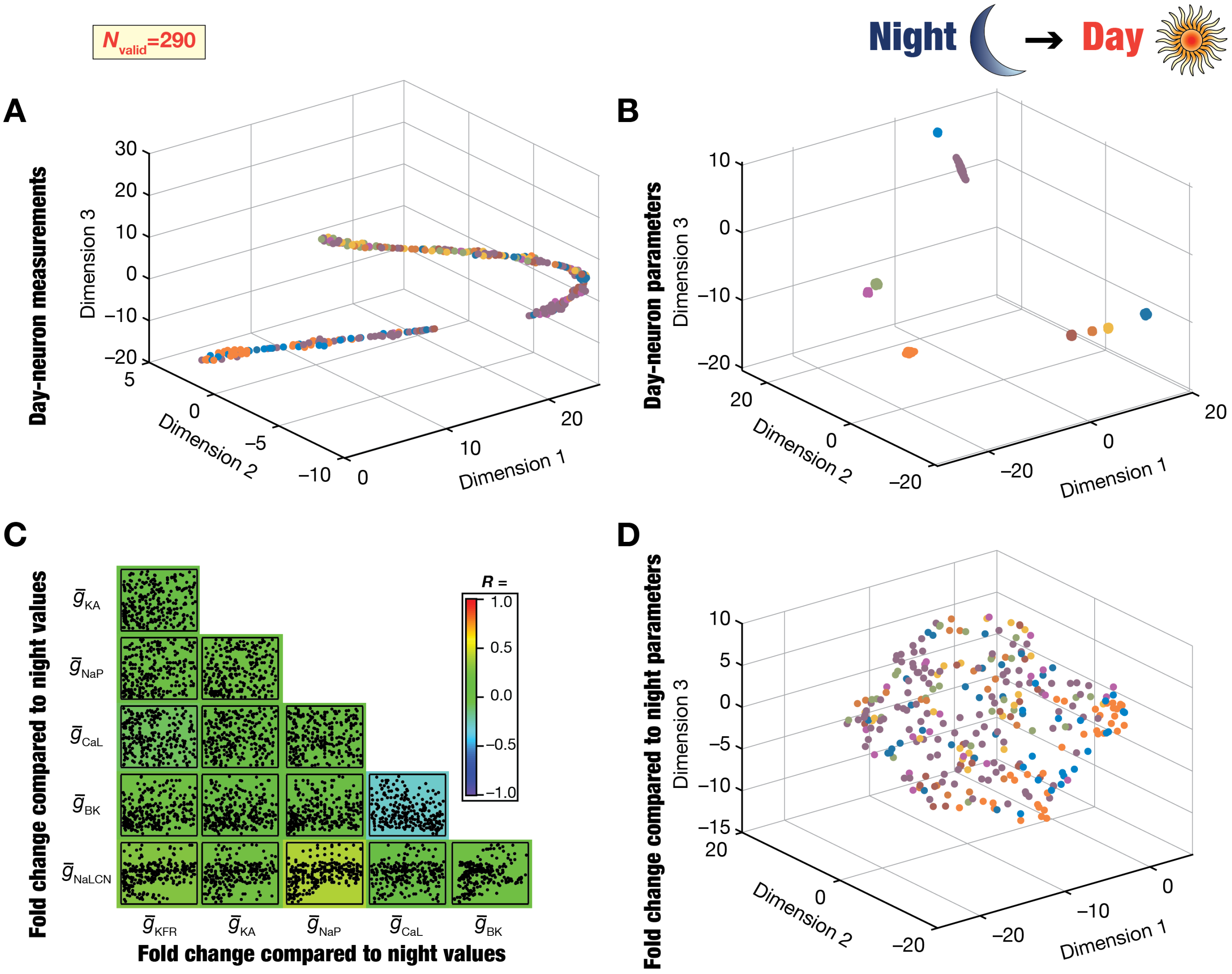

**Supplementary Figure S11. Analyses involving dimensionality reduction of fold-changes in ion-channel conductances involved in night-to-day transitions revealed the absence of plasticity manifolds from a *de novo* population of night-like neurons.** (*A*) Representation of the 9 measurements from all 290 day-like neurons on a reduced 3-dimensional space obtained through *t*-SNE. Different colors represent the 9 distinct night-like neurons from which the day-like models were obtained (shown in Supplementary Fig. S9). The absence of clustering based on the night-like neuron implies that day-like neurons with distinct origin may show similar measurement phenotypes. (*B*) Representation of the 13 parameters from all 290 night-like neurons on a reduced 3-dimensional space obtained through *t*-SNE. Different colors represent the 9 distinct night-like neurons from which the day-like models were obtained (shown in Supplementary Fig. S9). The parameters associated with the day-like neurons formed distinct clusters based on night-like neuron where they transitioned from. (*C*) Scatter-plot matrix showing pairwise relationships between the fold changes in the six ion-channel conductances that underwent plasticity to yield the 290 valid day-like neurons from the 9 night-like neurons. The background color represents the value of the Pearson correlation coefficient between the different fold changes, and indicate weak pairwise correlations across all pairs. (*D*) Representation of the fold changes in the 6 ion-channel conductances that yielded the 290 day-like neurons, with reference to conductance values in their respective night-like counterparts. Fold changes are shown on a reduced 3-dimensional space obtained through *t*-SNE. Different colors represent the 9 distinct night-like neurons from which the day-like models were obtained (ion channel distributions in night-like neurons shown in Supplementary Fig. S9). Note the absence of clustering based on night-neuron colors and the absence of plasticity manifolds defining the night-to-day transitions from a *de novo* population of night neurons.
